# YAP/TEAD4-regulated placental alkaline phosphatases ALPP and ALPPL2 are immunosuppressive ectonucleotidases modulated by MAPK inhibitors

**DOI:** 10.1101/2025.10.15.682438

**Authors:** Jacob L Martin, Bruce Huang, Tobias Ehrenberger, Ahmed R Hasan, Michelle Dow, Weng Hua Khoo, Romain Pacaud, Essam Metwally, Katherine Porth, Gukhan Kim, Sawyer McKenna, Cindy Li, Nicholas Ho, Khoa Nguyen, Spencer Hahn, Barbara Rath, Karthik Sathiyamoorthy, Samantha Shu Wen Ho, Peng Yang, Aleksandra Olow, Svetlana Sadekova, David Bauché

## Abstract

Placental alkaline phosphatases ALPP and ALPPL2 (ALPP/L2) are promising clinical targets for therapies such as antibody-drug conjugates (ADCs). However, their regulation and biological functions remain unclear. Here, we identify the YAP/TEAD4 signaling axis as the primary regulator of ALPP/L2 expression in tumor cells. Importantly, we demonstrate that inhibitors targeting KRAS, MEK, and ERK upregulate ALPP/L2 expression *in vitro* and *in vivo*, thereby enhancing the efficacy of antibody-drug conjugates (ADCs) when used in combination. Moreover, our work reveals a novel immunomodulatory function of ALPP/L2 as cell surface ectonucleotidases that hydrolyze extracellular ATP, ADP, and AMP to adenosine, promoting regulatory T cell infiltration into the tumor microenvironment. Together, these findings reveal a biological function of ALPP/L2 and identify druggable regulators to enhance ALPP/L2-targeted therapies.

**SIGNIFICANCE:** This study uncovers the YAP/TEAD pathway as a positive regulator of the clinically relevant tumor-associated antigens ALPP and ALPPL2. Kras, MEK, or ERK inhibitors enhance ALPP/L2 expression in tumor cells, thereby increasing the efficacy of ALPP/L2-targeted antibody-drug conjugates. These findings offer a promising combination strategy to overcome resistance to ALPP/L2-based therapies caused by low target expression.

## INTRODUCTION

Placental alkaline phosphatases ALPP and ALPPL2 (herein referred to as ALPP/L2) have recently gained attention as promising clinical targets for cell-based immunotherapies and targeted modalities such as antibody-drug conjugates (ADCs) and cell engagers (^1–4^, NCT05229900). ALPP/L2 are glycosylphosphatidylinositol (GPI)-anchored metalloenzymes predominantly expressed on the surface of normal placental and cervical tissues but overexpressed in multiple tumor types, including ovarian, endometrial, testicular, pancreatic, gastric, bladder, colorectal, and non-small cell lung cancers ^5–7^, making them attractive candidates for selective therapeutic intervention. In a phase 1 trial with a limited number of patients, objective tumor regression was observed in two of three ovarian cancer patients treated with TCA-101, an ALPP/L2-targeting chimeric antigen receptor (CAR)-T cell therapy ^8^ (NCT07108140), highlighting the potential of ALPP/L2 as therapeutic targets. Although these tumor-associated antigens were identified more than fifty years ago, the molecular mechanisms governing ALPP/L2 expression and their functional roles within the tumor microenvironment remain largely unexplored.

Understanding the druggable regulatory pathways that control ALPP/L2 expression is critical for optimizing therapeutic strategies that exploit these targets. In the current study, we identified TEAD4 and NF2, two proteins involved in the Hippo pathway ^9,10^, as regulators of ALPP/L2 expression in human cancer cell lines. Recent studies have shown that KRAS G12C inhibitors, such as sotorasib, enhance YAP/TEAD transcriptional activity and that the YAP/TAZ/TEAD pathway mediates resistance to Kras inhibitors ^11–13^. Mechanistically, Adachi et al. proposed that Kras inhibitors prevent Scrib-mediated inhibition of YAP, allowing YAP translocation to the nucleus ^11^. MEK and ERK, two proteins downstream of Kras, also inhibit YAP/TEAD activity in YAP/TEAD-dependent cells ^14^. Moreover, the impact of clinically approved inhibitors targeting the Kras-MEK-ERK signaling axis on ALPP/L2 expression and subsequent therapeutic efficacy remains unknown.

Extracellular adenosine is a metabolic and immunosuppressive molecule generated by the stepwise conversion of ATP, ADP, and AMP. Numerous studies have shown that CD39 and CD73, expressed by immune and non-immune cells, cooperate to convert extracellular ATP to adenosine ^15,16^. Adenosine binding to its receptors promotes infiltration and function of immunosuppressive cells such as regulatory T cells (Tregs) and myeloid cells, while inhibiting anti-tumoral T and NK cells ^17,18^. The tissue non-specific alkaline phosphatase (TNAP), another alkaline phosphatase family member sharing 54% and 56% identity with ALPP and ALPPL2, respectively, has been shown to hydrolyze ATP stepwise to adenosine ^19^. Therefore, we hypothesized that ALPP/L2 promote tumor progression by creating an immunosuppressive tumor microenvironment via conversion of extracellular ATP to adenosine.

Here, we identify the YAP/TEAD4 pathway as a key regulator of ALPP/L2 expression in tumor cells and demonstrate that inhibition of the KRAS-MEK-ERK pathway upregulates ALPP/L2 both *in vitro* and *in vivo*, including in human patients. We further show that combining these small-molecule inhibitors with ALPP/L2-targeted ADCs significantly enhances tumor cell killing. Finally, we reveal that ALPP/L2 ectonucleotidase activity facilitates Treg infiltration into tumors, highlighting a dual role for ALPP/L2 in tumor biology and immunomodulation.

## RESULTS

### Identification of regulators of ALPP/L2 using a genome-wide CRISPR screen

The human alkaline phosphatases ALPP and ALPPL2 (ALPP/L2) are normally expressed in the placenta and cervix and are overexpressed in multiple cancer types, including ovarian, endometrial, pancreatic (PDAC), and non-small cell lung cancers (NSCLC) ^6^. Although ALPP and ALPG (gene encoding for ALPPL2) are distinct genes, they are co-expressed in ovarian and endometrial cancer samples as well as in human cancer cell lines (**Figure S1**). We first evaluated ALPP/L2 expression in dissociated tumor cells from ovarian cancer patients using flow cytometry and immunohistochemistry (**Figure 1A, D**). We found that ALPP/L2 expression is highly heterogeneous (**Figure 1B, E**) and generally low, with an average of approximately 5,000 ALPP/L2 molecules per tumor cell (herein referred to as copy number) (**Figure 1C**). Intratumoral heterogeneity is thought to limit the efficacy of targeted modalities such as antibody-drug conjugates (ADCs), chimeric antigen receptor T cells (CAR-T), or T-cell engagers (TCEs) ^20–22^. Therefore, our goal was to identify common druggable regulators of ALPP and ALPPL2 expression in tumor cells. To this end, we performed a genome-wide CRISPR screen in ALPP KO and ALPPL2 KO SW837, a human cancer cell line that is easily transfectable and expresses comparable levels of ALPP and ALPPL2 (**Figure 1F**). Several genes were identified as common negative or positive regulators (**Figure 1G**). As expected, along with ALPG and ALPP, genes involved in glycosylphosphatidylinositol (GPI) anchor biosynthesis ranked among the top positive regulators of ALPP/L2 (**Figure 1H, Figure S2**). Notably, TEAD4, a placental-enriched transcription factor commonly amplified in indications in which ALPP/L2 is expressed —such as ovarian, NSCLC or PDAC cancer— and involved in the Hippo pathway^9,23^ (**Figure S3**), ranked among the top significant positive regulators, whereas NF2, a negative regulator of the Hippo pathway^10,24^, ranked as a top significant negative regulator of ALPP/L2 (**Figure 1G, I**). Altogether, our data suggest that ALPP/L2 are heterogeneous tumor-associated antigens regulated by the YAP/TAZ/TEAD pathway.

**Figure 1:**
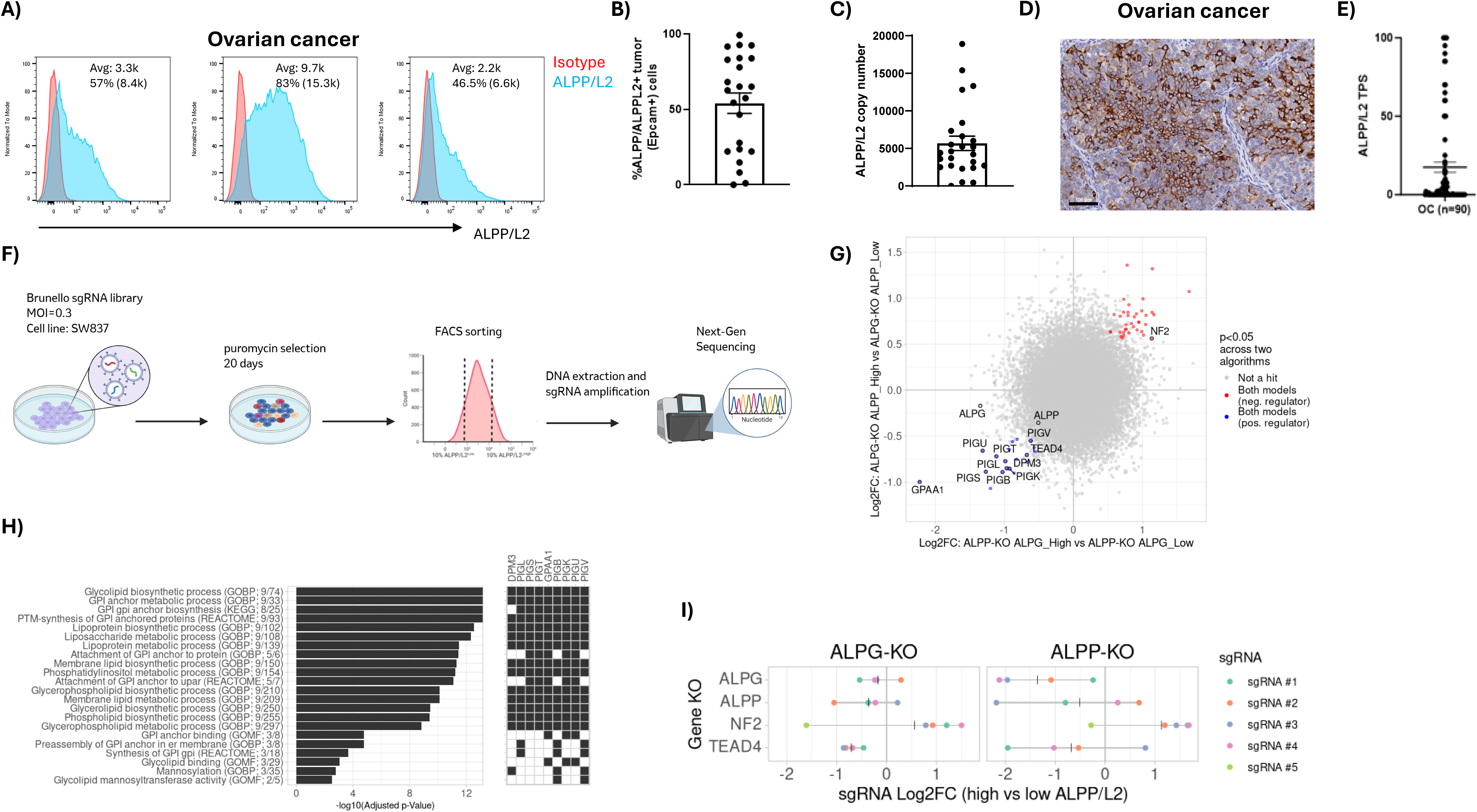
Identification of regulators of ALPP/L2 using a genome-wide CRISPR screen. **(A)** Cell surface expression of ALPP/L2 on human primary ovarian tumor cells. **(B)** Percentage of ALPP/L2-positive cells among EpCAM-positive cells. **(C)** ALPP/L2 copy number per tumor cell. n = 25 samples from ovarian cancer patients. **(D)** Representative image of ALPP/L2 expression in ovarian cancer by immunohistochemistry. **(E)** Quantification of ALPP/L2 expression shown as the ALPP/L2 Tumor Proportion Score (TPS). n = 90 samples. **(F)** Schematic of the genome-wide CRISPR screen performed on ALPPL2 knockout (ALPPL2KO/ALPGKO) or ALPP knockout (ALPPKO) SW837 cells to identify regulators of ALPP and ALPPL2, respectively. Created using BioRender (https://BioRender.com). **(G)** Positive (blue dots) and negative (red dots) regulators of ALPPL2 in SW837 cell lines, ranked based on the average fold changes of their targeting sgRNAs and p-values. Labeled genes are either controls (ALPP, ALPG) or other genes of interest (TEAD4, NF2, and GPI-associated). **(H)** Overrepresentation analysis of the positive regulators of ALPP and ALPPL2 in SW837 cells showing significantly enriched GO terms and gene sets (FDR-adjusted p value < 0.01). The left panel shows gene sets and GO term names alongside the number of hit genes and set sizes in parentheses. The right panel highlights which hit genes were associated with these gene sets. **(I)** Fold change of ALPP, ALPG, NF2, and TEAD4 in ALPGKO- or ALPPKO-high versus ALPGKO- or ALPPKO-low cells from individual sgRNAs. Colors distinguish the different sgRNAs for each gene to enable comparison across screens; vertical bars indicate averages across sgRNAs. Data are represented as mean ± SEM. Scale bar:100 µm

### TEAD4 induces ALPP/L2 expression in human cancer cell lines

To confirm TEAD4 and NF2 as regulators of ALPP/L2 in cancer cell lines, we generated TEAD4 and NF2 knockout (KO) SW837 cells. Using flow cytometry, western blot and RT-qPCR analysis, we showed that ALPP/L2 expression is significantly reduced in TEAD4 KO cells and significantly upregulated in NF2 KO cells (**Figure 2A-D**). Previous studies have identified TEAD4 as a transcription factor that recruits YAP/TAZ and binds to open chromatin or super-enhancer regions located more than 10 kb from transcription start sites ^25,26^. Interestingly, using available ChIP-seq data (GSE131687 ^27^, GSE44416 ^28^, GSE32465 ^29^), we found that both TEAD4 and YAP1 bind to both ALPP and ALPPL2 promoters in ALPP/L2-positive cell lines but not in negative ones.

**Figure 2:**
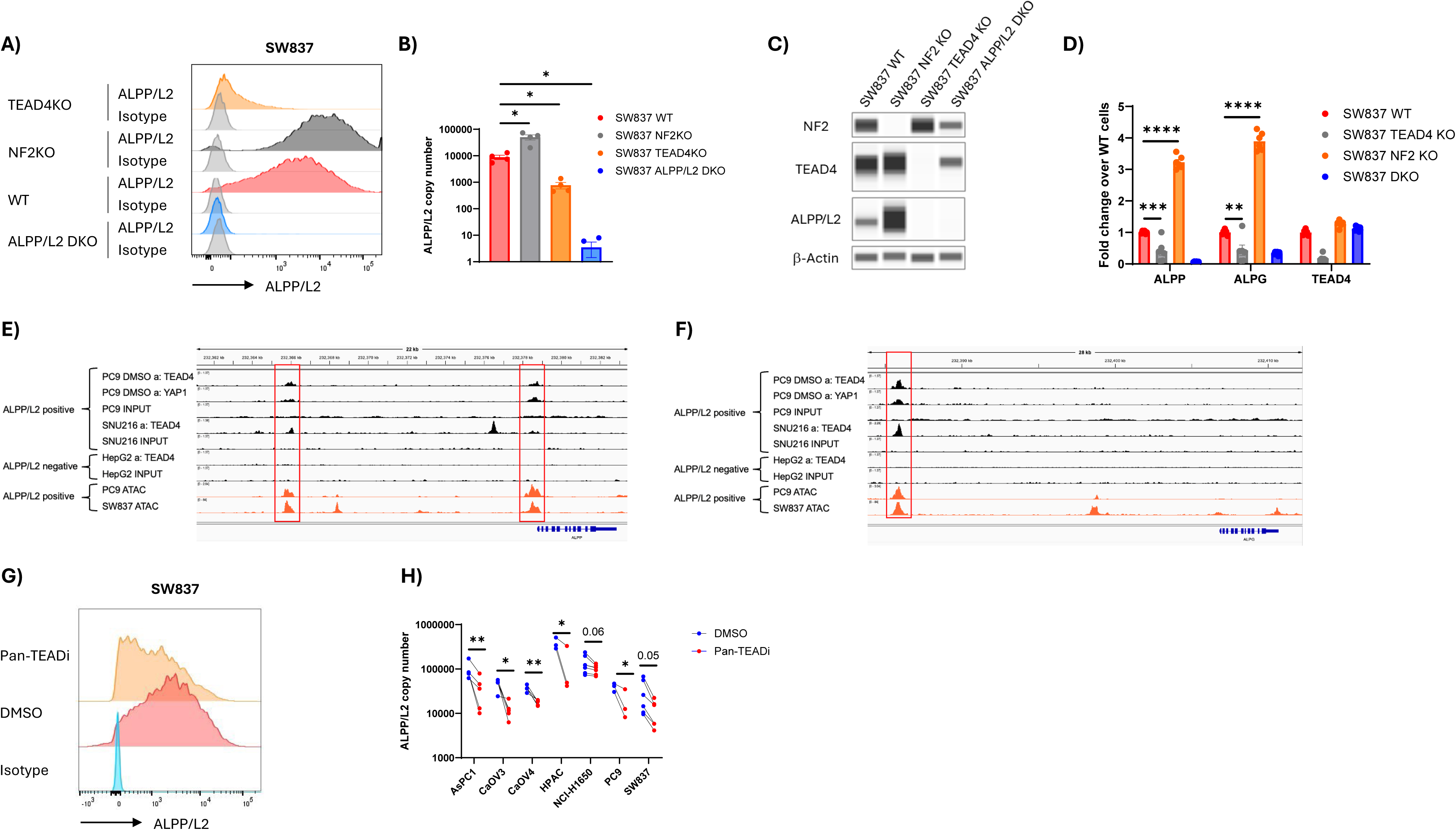
TEAD4 binds to ALPP and ALPPL2 promotes and regulates their expression. **(A)** Representative histogram of cell surface expression of ALPP/L2 on SW837 ALPP/L2 DKO, SW837 WT, SW837 TEAD4 KO, or SW837 NF2 KO cells evaluated by flow cytometry. **(B)** Quantification of ALPP/L2 copy number in the listed cell lines. **(C**) TEAD4, NF2, and ALPP/L2 expression was evaluated by western blot. **(D**) TEAD4 ALPP and ALPG expression was measured by RT-qPCR. **(E-F**) ChIP-seq peaks demonstrating binding of TEAD4 and/or YAP1 in ALPP (**E**) and (**F**) ALPG promoters in ALPP/L2 positive PC9 and SNU-216 cells but not ALPP/L2 negative HepG2 cells (black histograms). ChIP-seq data obtained from the ChIP-atlas (https://chip-atlas.org/). TEAD4 binds to open regions as shown by ATAC-seq (orange histograms) (GSM7112169). **(G**) Representative histogram of cell surface expression of ALPP/L2 of SW837 cells treated with a pan-TEAD inhibitor at 10 μM for 3 days. **(H**) ALPP/L2 copy number of the listed human cancer cell lines treated with a Pan-TEADi at 10uM for 3 days. Data are represented as mean ± SEM. **(A-D, G-H**) n=2-6, **(E-F) n=1** independent experiments. **(A-B**) One way ANOVA * p<0.05, ** p<0.01, *** p<0.001, **** p<0.001; **(G**) Paired t-test * p<0.05, ** p<0.01

Specifically, there is one binding site characterized by the consensus sequence CATTCCA/T ^30^ in the ALPPL2 promoter (∼20 kb from the start site) and two binding sites in the ALPP promoter (one near the start site and another ∼13 kb from the start site) **(Figure 2E-F)**. As previously reported, TEAD4 binding occurs only in the open chromatin regions of ALPP/L2-positive cells, as shown by ATAC-seq **(Figure 2E-F)**. Next, we performed western blot analysis of cytoplasmic and nuclear fractions from human cancer cell lines expressing various levels of ALPP/L2. Although we did not observe a clear correlation between TEAD4 nuclear localization and ALPP/L2 expression, we found that ALPP/L2 expression correlates with chromatin accessibility at the TEAD4 binding sites identified by ChIP-seq, suggesting epigenetic regulation in ALPP/L2-negative cancer cells **(Figure S4).**

We then assessed whether pharmacological inhibition of the YAP/TEAD pathway modulates ALPP/L2 expression in human cancer cell lines by analyzing an RNA-sequencing dataset (GSE229071) ^13^. GNE-7883, a pan-TEAD inhibitor that blocks YAP/TAZ association with TEAD and reduces chromatin accessibility, significantly reduces ALPP—but not ALPG—expression in OVCAR8 and HCC1576 cell lines (**Figure S5A-B**). We further confirmed that a pan-TEADi induces a dose-dependent downregulation of ALPP/L2 in human cancer cell lines from multiple indications (**Figure 2G-H, Figure S5C)**. Our data confirm that TEAD4 is a positive regulator of ALPP/L2 expression and that TEAD inhibitors suppress its expression *in vitro*.

To evaluate the functional impact of ALPP/L2 regulation *in vitro*, we treated SW837 cells expressing various levels of ALPP/L2—from 0 to 800,000 copies per cell—with an ALPP/L2- MMAE conjugate (**Figure S6A**). We demonstrated a positive correlation between ALPP/L2 expression and *in vitro* potency, as shown by IC50 values (**Figure S6B-C**). Notably, ALPP/L2- MMAE-mediated killing was not observed in TEAD4 KO cells, while the antibody-drug conjugate (ADC) kills NF2 KO cells at sub-nanomolar concentrations (**Figure S6B**). Our data suggest that modulating ALPP/L2 expression in tumors can enhance ADC-mediated killing.

### Kras/MEK/ERK inhibitors enhance TEAD4-mediated ALPP/L2 expression in vitro and in vivo

Recent studies have highlighted the role of Kras in regulating the YAP/TAZ/TEAD pathway in preclinical models and patients. Notably, the YAP/TAZ/TEAD pathway mediates resistance to Kras inhibitors such as the FDA-approved Sotorasib and MRTX1133^13,31^. Because the YAP/TAZ/TEAD pathway is activated in Kras inhibitor–acquired resistant cells, we hypothesized that ALPP/L2 expression can be enhanced by Kras inhibitors. To test this, we generated Kras G12C inhibitor (G12Ci)-acquired resistant lung cancer cell line NCI-H358, here named NCI-H358R (**Figure S7A**). Using RNA-seq, we found that ALPP and ALPG are significantly upregulated in NCI-H358R cells (**Figure 3A**). Similarly, RNA-seq data from a previous study reveal that ALPP and ALPG are upregulated in AMG510-acquired resistant NCI-H23 cell lines compared to DMSO-treated cells (**Figure S7B**, GSE229071 ^13^). We further confirmed that ALPP/L2 is significantly overexpressed at the cell surface of NCI-H358R cells compared to DMSO-treated NCI-H358 cells by flow cytometry (**Figure 3B**). In line with our previous observations (**Figure S6**), NCI-H358R cells were more sensitive to ALPP/L2-MMAE–mediated killing than DMSO-treated NCI-H358 cells *in vitro* (**Figure 3C**), while both lines were equally sensitive to free MMAE (data not shown).

**Figure 3:**
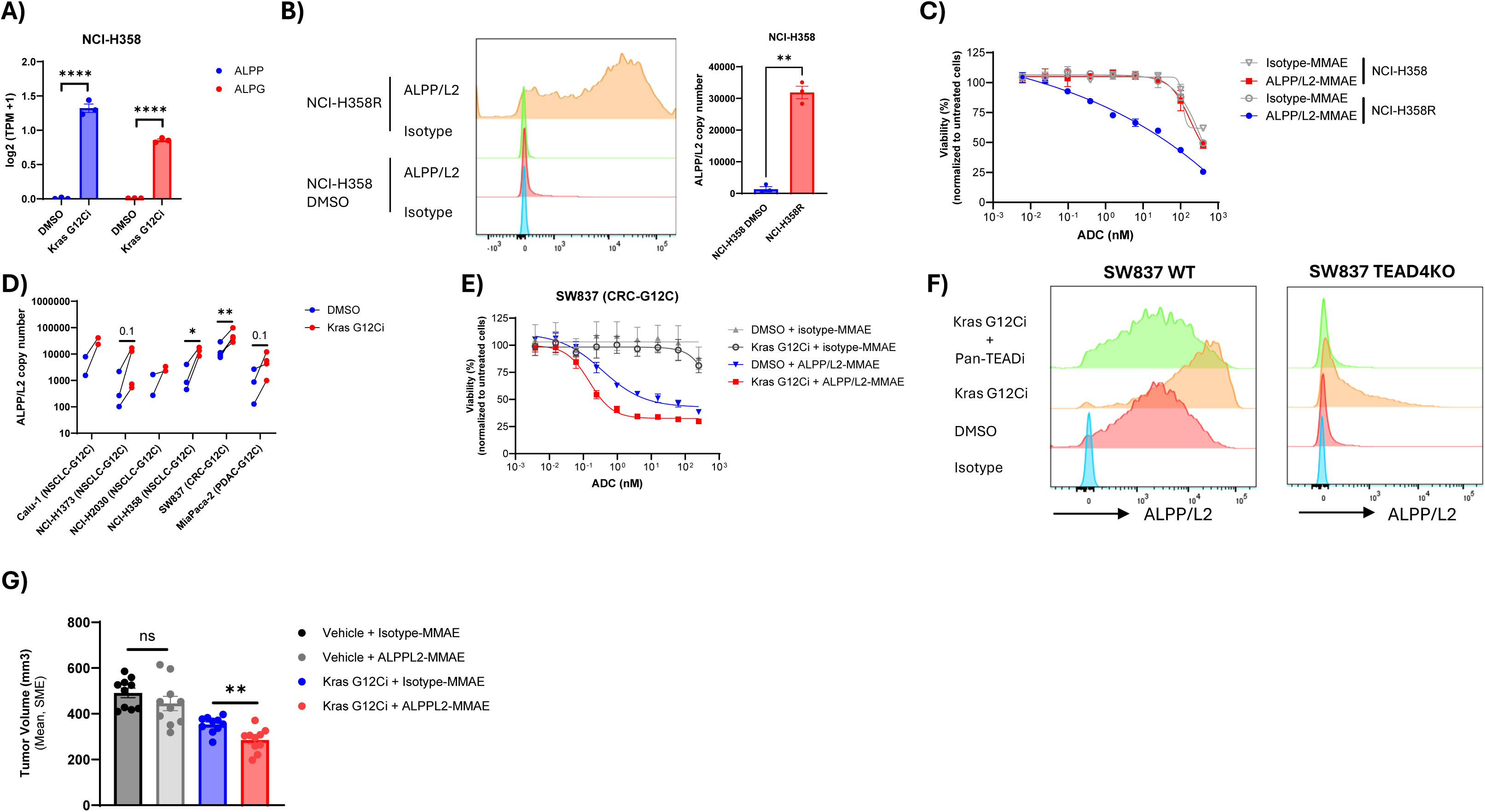
Kras G12Ci treatment upregulates ALPP/L2 expression *in vitro* and *in vivo* via TEAD4. **(A)** ALPP and ALPG expression in Kras G12Ci-acquired resistant NCI-H358 cells (NCI-H358R) compared to DMSO-treated NCI-H358 cells, measured by RNA-seq. **(B)** Representative histogram and quantification of ALPP/L2 cell surface expression on NCI-H358 DMSO or NCI-H358R cells *in vitro*. **(C)** Cell viability of NCI-H358 DMSO or NCI-H358R cells treated with different concentrations of ALPP/L2-MMAE or isotype-MMAE conjugates for 6 days. **(D)** ALPP/L2 copy number in human cancer cell lines harboring a KRAS G12C mutation treated with 62.5 nM Kras G12Ci for three days. NCI-H322 (KRAS WT) was used as a negative control. **(E)** Cell viability of SW837 cells pre-treated with Kras G12Ci (62.5 nM) or DMSO for two days, followed by treatment with different concentrations of ALPP/L2-MMAE or isotype-MMAE conjugates for 6 days. **(F)** Representative histogram and quantification of ALPP/L2 cell surface expression on SW837 WT (left) or SW837 TEAD4KO (right) cells treated with 62.5 nM Kras G12Ci, 10 µM Pan-TEADi, or the combination for three days *in vitro*. **(G)** SW837-bearing NSG mice were dosed daily with Kras G12Ci (10 mpk) or vehicle control for 8 doses. On day 2, mice received a single dose of ALPP/L2-MMAE or isotype-MMAE. Tumor volume was measured at day 18 post-treatment. N = 10 mice per group, 1 independent experiment. Data are represented as mean ± SEM. (A-F) n = 3–6 independent experiments. Statistical analysis: (A, G) unpaired t-test, **** p < 0.0001; (B, D) paired t-test, ns = not significant, * p < 0.05, ** p < 0.01.

Next, we evaluated whether short-term inhibition of Kras G12C also upregulates ALPP/L2 via the YAP/TEAD pathway. Treatment of human cell lines from various cancer types—including NSCLC, PDAC, and CRC—harboring Kras G12C mutations with a Kras G12C inhibitor significantly increased ALPP/L2 surface expression (**Figure 3D, Figure S7C**). Consequently, pre-treatment with Kras G12Ci sensitized SW837 cells to ALPP/L2-MMAE–mediated killing *in vitro* (**Figure 3E**) while sensitivity to free MMAE remained unchanged (data not shown). Mechanistically, Kras G12Ci-mediated upregulation of ALPP/L2 was mainly driven by TEAD4, as the Kras G12Ci failed to significantly upregulate ALPP/L2 in TEAD4 knockout cells (**Figure 3F**). Notably, other TEAD family members appear to partially control ALPP/L2 expression, as treatment with a pan-TEAD inhibitor completely abrogated Kras G12Ci-mediated residual upregulation in TEAD4 knockout cells (**Figure 3F**). Finally, we evaluated the impact of Kras G12Ci combined with ALPP/L2-MMAE *in vivo*. Strikingly, Kras G12Ci pre-treatment for two days boosted ALPP/L2 surface expression on SW837 cells and significantly enhanced ALPP/L2- MMAE–mediated antitumor efficacy *in vivo* (**Figure 3G, Figure S7D**).

Kras is the most frequently mutated oncogene in cancer ^32^. While Kras G12C was the first mutation to be successfully targeted, other mutations such as G12D or G12V are more prevalent, especially in CRC and PDAC ^32,33^. To test whether ALPP/L2 upregulation is specific to the Kras G12Ci or extends to other clinical Kras inhibitors, we treated human cancer cell lines harboring various Kras mutations—including G12D, G12V, and G13D—as well as a KRAS wild-type ovarian cancer cell line with a pan-Ras inhibitor (pan-Rasi). Similar to our previous observation, the Pan-Rasi induces a dose-dependent upregulation of ALPP/L2 in cancer cell lines *in vitro* (**Figure 4A-B**). Because ERK and MEK are downstream of Kras and are druggable targets, we next evaluated two small-molecule inhibitors, targeting MEK and ERK, respectively ^34,35^. Consistent with previous findings, MEK inhibitor (MEKi) and ERK inhibitor (ERKi) induced dose-dependent upregulation of ALPP/L2 in multiple human cancer cell lines (**Figure 4C-D, Figure S8A-F**), as early as 24 hours post-treatment (**Figure S8G**). Altogether, our data suggest that targeting the KRAS/MEK/ERK pathway with small-molecule inhibitors can enhance ALPP/L2-targeted ADC efficacy by significantly upregulating ALPP/L2 expression in tumor cells (**Figure 4E)**.

**Figure 4:**
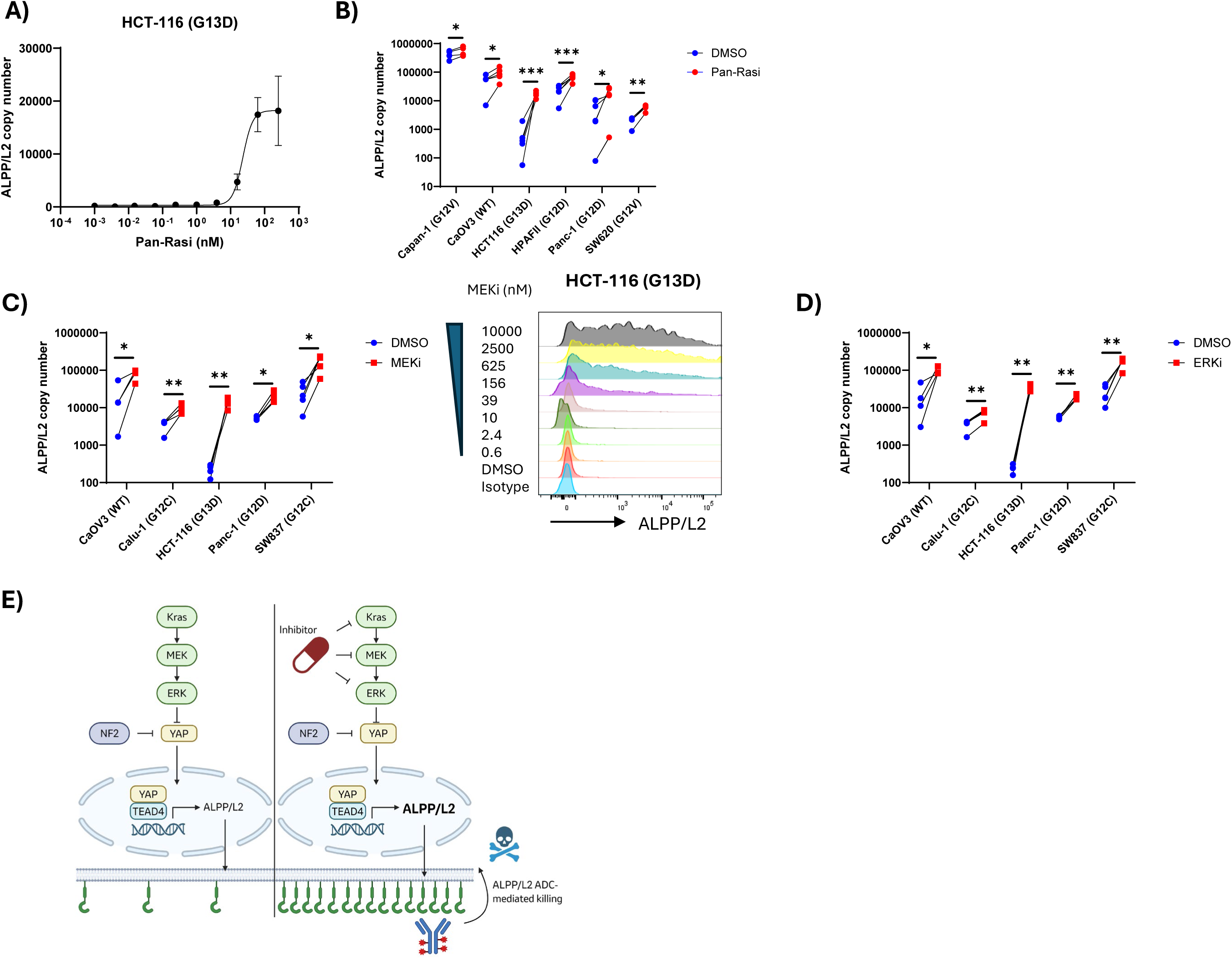
The Kras/MEK/ERK pathway regulates ALPP/L2 expression *in vitro* and *in vivo*. **(A)** Dose-dependent upregulation of cell surface ALPP/L2 on HCT-116 cells treated with Pan-Rasi. **(B)** ALPP/L2 copy number in Kras wild-type (WT) or Kras mutant human cancer cell lines treated with 15.6 nM (or 62.5 nM for HCT116) Pan-Rasi for three days. **(C)** ALPP/L2 copy number (left) and representative histograms (right) of KRAS WT or Kras mutant human cancer cell lines treated with 2.5 µM MEKi for three days. **(D)** ALPP/L2 copy number of Kras WT or Kras mutant human cancer cell lines treated with 2.5 µM ERKi for three days. **(E)** Schematic representing the regulation of ALPP/L2 expression by YAP/TEAD4. Kras/MEK/ERK inhibitors enhance ALPP/L2 expression and ALPP/L2 ADC-mediated killing. Schematic created using BioRender (https://BioRender.com). Data are represented as mean ± SEM. (A–D) n = 3–6 independent experiments. Statistical analysis: (B–D) paired t-test, ns = not significant, * p < 0.05, ** p < 0.01, *** p < 0.001.

### ALPP/L2 are ectonucleotidase enzymes promoting Treg infiltration in human and mouse tumors

Although ALPP/L2 expression in tumors has been extensively reported, the biological functions of these proteins remain largely unknown. Consistent with a previous report, we found that high ALPP/L2 expression correlates with poor prognosis in multiple tumor types (**Figure S9A-C**) ^36^, suggesting a pro-tumoral role for these enzymes. Interestingly, TEAD4 expression correlates with Treg cell infiltration during skin inflammation^37^. In addition, NF2 loss-of-function or acquired resistance to sotorasib also correlates with an immunosuppressive microenvironment characterized by increased tumor-infiltrating Treg cells ^38,39^. Since ALPP/L2 are GPI-anchored proteins lacking intracellular signaling domains and are regulated by TEAD4/NF2 and Kras inhibitors, we hypothesized that these enzymes promote tumor growth by recruiting Treg cells.

We found that the proportion of ALPP/L2-expressing tumor cells positively correlates with the proportion of Treg cells in human ovarian and endometrial tumor samples (**Figure 5A-B)**. We further confirmed in mouse syngeneic models that overexpression of human ALPPL2 in MC38 and MB49 tumor cells leads to increased tumor-infiltrating Treg cells *in vivo* compared to empty vector controls (**Figure 5C, Figure S10)**. Notably, although the proportion of CD4+ T cells increased, we observed no significant differences in infiltration of other lymphoid cells, such as CD8+ T cells, NK cells, or B cells, or myeloid cells, such as gMDSC, mMDSC, or Macrophages (**Figure S11)**.

**Figure 5:**
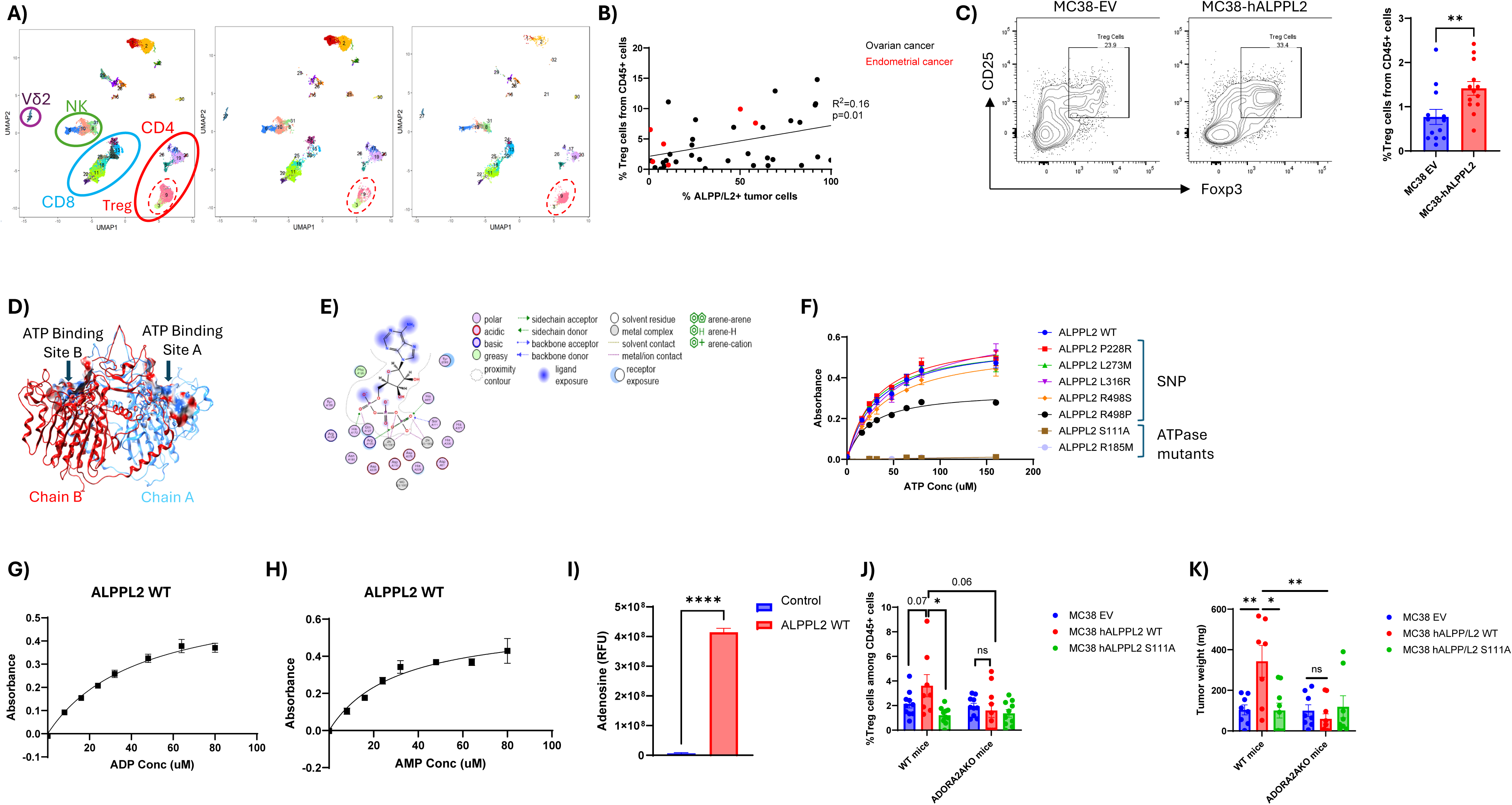
ALPP/L2 are ectonucleotidase enzymes promoting Treg infiltration in human and mouse tumors. **(A)** UMAP plots showing T and NK cells subsets (gated on live CD45+ CD19-CD11b-CD11c-cells) from ALPP low (<1500 copies/tumor cell) and ALPP/L2 high (>1500 copies/tumor cell) ovarian cancer samples. n=3-4 samples. **(B)** Correlation of ALPP/L2 expression and Treg cells infiltration in human ovarian (n=32) and endometrial (n=6) tumor cells. **(C)** Representative flow cytometric analysis of Treg cells (live CD45+CD3+CD4+Foxp3+CD25+) (left) and quantification (right) in MC38-EV and MC38-hALPPL2 tumors, day 21 post-implant. **(D)** Crystal structure of the human placental alkaline phosphatase dimer (PDB 1EW2), highlighting the ATP binding sites. **(E)** Simulation of ATP binding to the binding sites. **(F-H)** Phosphatase activity of ALPPL2 WT, SNPs, S111A, and R185M mutants using malachite green assay, **(F)** ATP, **(G)** ADP, **(H)** AMP was used as substrate. **(I)** Hydrolysis of ATP to adenosine by ALPPL2 WT recombinant protein. **(J)** Quantification of tumor-infiltrating Treg cells (live CD45+CD3+CD4+Foxp3+CD25+) in WT or ADORA2KO mice implanted with MC38-EV, MC38-hALPPL2, and MC38-hALPPL2 S111A tumors, day 21 post-implant. **(K)** Weight of MC38-EV, MC38-hALPPL2, and MC38-hALPPL2 S111A tumors implanted in WT or ADORA2KO mice, at day 21 post-implant. Data are represented as mean ± SEM. **(B**-**C, F-K**) n=2 independent experiments. **(B, C, J, K)** Unpaired t-test ns=non-significant, * p<0.05, ** p<0.01. **(I)** Paired t-test **** p<0.01

To understand how ALPP/L2 modulates the tumor microenvironment, we examined the crystal structure of ALPP. Several pockets and metal ions (Zn^2+^, and Mg^2+^) were identified in the crystals. These ions, in conjunction with an analysis of potential binding sites, helped identify a potential substrate for other alkaline phosphatases ^19^ (**Figure 5D)**. Docking of ATP to the identified pocket suggested that Serine 111 (S111) and Arg 185 (R185) are key anchoring amino acids involved in the catalytic domain (**Figure 5E)**. We next evaluated the impact of single amino acid mutations on the enzymatic function of ALPP and ALPPL2. Both wild-type ALPP and ALPPL2 proteins hydrolyze ATP (**Figure 5F, Figure S12A),** ADP (**Figure 5G, Figure S12B),** and AMP (**Figure 5H, Figure S12C),** leading to the production of adenosine **(Figure 5I)** *in vitro*, suggesting a stepwise conversion of ATP to adenosine and a redundant role for both ALPP and ALPPL2. In contrast, mutations of S111 or R185 abrogate ALPPL2-mediated ATP hydrolysis (**Figure 5F),** validating ALPP/L2 as novel tumor-associated ectonucleotidases.

Finally, we investigated whether ALPPL2-mediated Treg infiltration is driven by adenosine *in vivo*. We generated human ALPPL2 S111A-expressing MC38 cells (MC38-hALPPL2 S111A) with expression levels matching those of MC38-hALPPL2 wild-type (WT) cells (**Figure S12D)** and confirmed that MC38-hALPPL2 S111A cells lack the ability to hydrolyze ATP, ADP, and AMP *in vitro* (**Figure S12E).** We then implanted these tumor cells into wild-type (WT) and ADORA2A knockout (ADORA2AKO) mice — an adenosine receptor expressed on immune cells, including Treg cells ^40,41^— to assess the role of adenosine signaling in Treg infiltration. Confirming our hypothesis, increased Treg infiltration was not observed in tumors harboring the S111A mutation or when MC38-hALPPL2 WT cells were implanted in ADORA2AKO mice (**Figure 5J)**. Consequently, MC38-hALPPL2 WT tumors were significantly larger in WT mice than in ADORA2KO mice (**Figure 5K)**. Altogether, our data demonstrate that ALPP/L2 are cell surface ectonucleotidases that recruit Treg cells into tumors via adenosine signaling.

## DISCUSSION

The regulation and biological function of the emerging clinical tumor-associated antigens ALPP and ALPPL2 remain poorly understood. In this work, we performed a genome-wide CRISPR screen that identified the YAP/TEAD pathway—particularly TEAD4—as the main positive regulator of ALPP/L2 expression in tumor cells. Recent studies have shown that YAP/TEAD confers resistance to Kras inhibitors, suggesting that Kras inhibitors can boost ALPP/L2 expression in tumor cells. Indeed, we demonstrated that KRAS, MEK, and ERK inhibitors enhance ALPP/L2 expression in human cancer cell lines via TEAD4. Furthermore, combining Kras G12C inhibitors—and potentially any Kras, MEK, or ERK inhibitors—with ALPP/L2-targeted ADCs enhances tumor cell killing *in vitro* and *in vivo*. This finding has significant clinical implications as ALPP/L2-targeting therapies are currently being developed or tested in clinical trials (NCT07108140, NCT05229900).

We found that TEAD4 is predominantly localized in the nucleus of both ALPP/L2-negative and - positive cell lines. Although TEAD4 nuclear localization does not correlate with ALPP/L2 expression, ATAC-seq data revealed that TEAD4 binding sites are inaccessible in the promoters of ALPP/L2-negative cells compared to positive cells, suggesting epigenetic regulation. While epigenetic drugs show promise for treating solid tumors ^42,43^, their efficacy remains limited. Based on our findings, combining epigenetic drugs with ALPP/L2-targeting therapies such as ADCs, CAR-T cells, or T-cell engagers could benefit cancer patients.

Although ALPP/L2 expressions in tumor cells have been extensively reported, their biological function was previously unknown. Here, we show that ALPP/L2 are cell surface ectonucleotidases that hydrolyze extracellular ATP, ADP, and AMP. Consequently, extracellular adenosine accumulates in the tumor microenvironment. Treg cells have been shown to accumulate in tumors from patients who developed resistance to sotorasib ^38^. Additionally, a recent study by Cole et al. demonstrated that Treg cells suppress MRTX1257 (a Kras G12C inhibitor)-mediated antitumor efficacy ^44^. While we showed that Kras inhibition enhances ALPP/L2 expression and that ALPPL2 promotes Treg infiltration in the tumor microenvironment, it remains to be demonstrated whether Kras inhibitor–induced Treg infiltration is driven by ALPP/L2 in patients.

In conclusion, we identified YAP/TEAD4 as positive regulators of ALPP/L2 expression in tumor cells. We demonstrated that small molecules targeting Kras, MEK, or ERK enhance ALPP/L2 expression and ALPP/L2-ADC–mediated killing *in vitro* and *in vivo*. Combining Kras inhibitors with ALPP/L2-targeted therapies holds great promise for improving cancer patient outcomes.

## MATERIALS AND METHODS

### Human samples

Frozen dissociated human tumor cells were from Discovery Life Science. Tumor surgical tissues were obtained from Seoul National University Hospital, Yonsei Cancer Centre, Taichung Veterans General Hospital where the collection protocol and informed consent have been approved by an Institutional Review Board (IRB)/ Ethics Committee (EC) in accordance with applicable country-specific regulatory requirements. Tissue microarrays from Ovarian cancer patients were obtained from TissueArray. Permission to perform this investigation was granted by the ethical committee, and written informed donor consent was obtained.

### Histology

Tumors were fixed in 10% neutral buffered formalin overnight, transferred to 70% ethanol, processed routinely, and embedded in paraffin. Human and rhesus monkey FFPE tissues were sectioned at 4 μm and then baked at 60°C for 45 minutes. Slides were subsequently deparaffinized using a Leica Stainer XL, subjected to heat-induced epitope retrieval, and washed with deionized water and TBST. Slides were loaded into an autostainer programmed as follows: 1) 3% H_2_O_2_ for 10 minutes, rinsed with 1x TBST; 2) Incubation with 1.5ug/ml of an anti-human/rhesus ALPP/L2 antibody (MS Validated antibodies, clone MSVa350R); 3) Incubation with the Envision Rabbit HRP (Dako) for 30 minutes, rinsed with 1x TBST; 4) Incubation with DAB solution (Dako) for 10 minutes, rinsed with 1x deionized water. Slides were then counterstained with Mayer’s Hematoxylin (Poly Scientific) using the Leica Stainer XL. Following counterstaining, slides were completely dried in a 60°C oven and cover-slipped with mounting medium (Dako) before microscopic examination. Slides were analyzed and scored by a certified pathologist.

### Fluorescence-activated cell sorting (FACS)

#### ALPP/L2 copy number quantification

Tumor samples and cancer cell lines were incubated with fixable viability dye eFluor450 (1:1000, ThermoFisher) and Human TrueStain FcX (Biolegend) for 25 minutes in 1x PBS. Then, cells were incubated with a saturating concentration (1 or 10 μg/mL) of phycoerythrin (PE)-conjugated ALPP/L2 antibody (Merck) or isotype control (clone QA16A12, Biolegend) for 30 minutes in stain buffer (BSA) (BD Pharmingen). Samples and Quantibrite PE beads (BD Biosciences) were run on a BD LSRFortessa™ flow cytometer (BD Biosciences) or Cytek Aurora (Cytek Biosciences). Data were analyzed using FlowJo software (FlowJo LLC). Mean Fluorescence Intensity (MFI) was converted to copy number (antibodies bound per cell).

### Mouse in vivo immunophenotyping

Single-cell suspensions from mouse tumors were incubated with fixable viability dye eFluor450 (ThermoFisher) and TruStain FcX (anti-mouse CD16/32) antibody (Clone 93, Biolegend) for 25 minutes in 1x PBS. Then, cells were incubated for 30 minutes in stain buffer (BSA) (BD pharmingen) with the following anti-mouse antibodies: CD45 BUV395 (Clone 30-F11, Biolegend); CD3 BV785 (Clone 17A2, Biolegend), CD4 PerCP (Clone GK1.5, Biolegend), CD8a APC (Clone 53-6.7, Biolegend), NK1.1 BUV737 (Clone PK136, BD Biosciences), CD25 APC/Cyanine7 (Clone PC61, Biolegend), CD19 FITC (Clone MB19-1, Biolegend) or CD19 PE (Clone 1D3/CD19, Biolegend), F4/80 BUV737 (Clone T45-2342, BD Biosciences), F4/80 BV785 (Clone M1/70, Biolegend), CD11c PerCP (Clone N418, Biolegend), I-A/I-E APC (Clone M5/114.15.2, Biolegend), Ly-6G PE (Clone 1A8, Biolegend), Ly-6C FITC (Clone HK1.4, Biolegend), CD24 APC/Cyanine7 (Clone M1/69, Biolegend). For intracellular Foxp3 staining, surface-stained cells were permeabilized, fixed with Foxp3 staining buffer set (Invitrogen) for 30 minutes on ice and then stained with Foxp3 PE or Alexa Fluor488 (Clone 150D, Biolegend) antibody for 30 minutes. Cells were washed twice and acquired on BD Fortessa flow cytometer (BD). Data were analyzed using FlowJo software (FlowJo).

### Human sample immunophenotyping

Frozen dissociated human tumor cells (Discovery Life Science) were incubated with fixable viability dye eFluor450 (ThermoFisher) and Human TruStain FcX (Biolegend) for 25 minutes in 1x PBS. Then, cells were incubated for 30 minutes in stain buffer (BSA) (BD pharmingen) with the following anti-human antibodies: Epcam APC (Clone 9C4, Biolegend), CD45 AlexaFluor 488 (Clone HI30, Biolegend), CD3 APC/Cyanine7 (Clone UCHT1, Biolegend), CD4 PerCP (Clone RPA-T4, Biolegend), CD8 PE/Cyanine7 (Clone SK1, Biolegend), CD25 PE (Clone BC96, Biolegend). Cells were washed twice and acquired on BD Fortessa flow cytometer (BD). Data were analyzed using FlowJo software (FlowJo).

Fresh human ovarian tumors were dissociated into single-cell suspensions as previously described^45^. Briefly, ovarian tumors were minced and dissociated in 100 U/mL Col IV (Worthington) + 400 U/mL DNase I (Worthington) at 37°C for 30 minutes with shaking. The cell suspension was filtered through a 70 µM cell strainer, washed with ice-cool PBS + 1% BSA, and spun at 450xG for 5 minutes at 4°C. Red blood cell lysis was performed using 1x RBC lysis buffer (ThermoFisher) at room temperature for 10 minutes. Cells were washed in ice-cool PBS + 1% BSA and spun at 450G for 5 minutes at 4 °C. Cell counts were performed manually using a C-Chip disposable hemacytometer (InCyto).

Dissociated ovarian tumors were stained for flow cytometry as previously described^45^. Briefly, cell suspensions were pelleted and stained with a viability dye, ViaDye Red Fixable Viability Dye (Cytek Biosciences) according to the manufacturer’s instructions. Next, cells were incubated with Human Fc Block (1:50, BD Biosciences) for 5 minutes at room temperature and incubated for 30 minutes in stain buffer (BSA) (BD Pharmingen) with the following anti-human antibodies: CD45 BUV395 (Clone H130, BD Biosciences), EPCAM BUV805 (Clone KS1/4, BD Biosciences) and (saturating concentration 1µg/mL) ALPP/L2 PE (Merck) or isotype control PE (Clone REA293, Miltenyi Biotec) in stain buffer (BSA). Cells were washed once in stain buffer, fixed in 1x Fix/Perm (BD Biosciences) for 30 minutes, and washed in 1x Perm/Wash (BD Biosciences). Cells were resuspended in stain buffer and acquired on the Cytek Aurora (Cytek Biosciences). Data were analysed using FlowJo (BD Biosciences).

### TCGA and GTEx analysis

For comparison of *ALPP* and *ALPG* expression between tumors and healthy tissues, a combined cohort of TCGA and GTEx RNA expression data was obtained from the UCSC Xena portal (RSEM tpm (n=19,131) UCSC Toil RNA-seq Recompute (UCSC Xena (xenabrowser.net)^46^.Tumor data was subset to primary disease samples only. Tumor and normal samples were paired by the provided primary site annotation. Data analysis and visualization were performed with R statistical programming language (4.4.2) and tidyverse (2.0.0).

### Cell lines

Human cancer cell lines HPAC, AsPC-1, CaOV3, CaOV4, NCI-H1650, PC9, NCI-H358, NCI- H322, NCI-H2122, Calu-1, NCI-H1373, NCI-H2030, MiaPaca-2, Capan-1, HCT-116, HPAF-II, SW837, Panc-1, SW620, and mouse cell lines MB49 and MC38 were purchased from the American Type Tissue Collection (ATCC). The cells were cultured in DMEM media (ThermoFisher) containing 10% heat-inactivated fetal calf serum (FCS) (GE Life Sciences), 100 U/ml penicillin (ThermoFisher), 100 µg/ml streptomycin (ThermoFisher), 1 % Non-Essential amino acids, 1% HEPES (4-(2-Hydroxyethyl)piperazine-1-ethane-sulfonic acid) 1% L-glutamine, and 1% Sodium Pyruvate (Gibco). All cells were cultured in a humidified incubator at 37 °C with 5% CO_2_. HPAC ALPP/L2 DKO, TEAD4 KO, and NF2 KO cells were generated by EditCo (formerly Synthego) using CRISPR technology. Briefly, parental WT cells were transfected with Cas9-sgRNA complex. For ALPP/L2 DKO, ALPPKO HPAC clone was then transfected with Cas9-ALPPL2 sgRNA complex, isolated as a single clone, expanded, and sequenced.

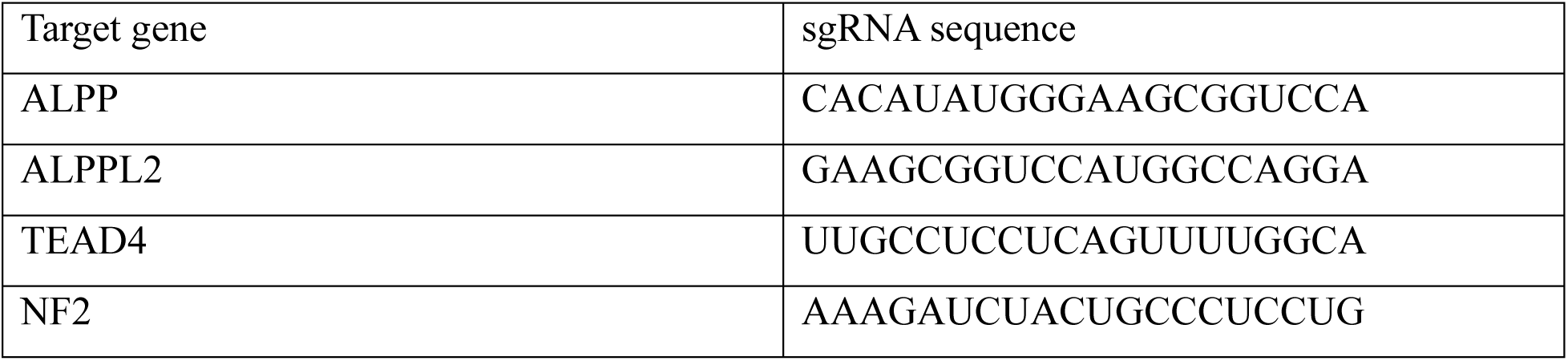

Human ALPPL2 WT or ALPPL2 S111A were cloned into a pLenti-EF1-BSD vector (Genscript). SW837, MC38, and MB49 were transfected with the lentiviral vectors (MOI=10) in the presence of 4 μg/ml of polybrene (Sigma-Aldrich). Cells were selected for at least two weeks with 8 µg/ml of blasticidin (Gibco).

### Genome-wide Pooled CRISPR screen

ALPPKO or ALPPL2KO SW837 cells were transduced with the lentiviral Brunello library at a multiplicity-of-infection (MOI) of 0.3 and polybrene at 8 µg/ml. After 16hours, the media was replaced. Cells were selected for 20 days with 2 μg/ml of puromycin (Gibco). At day 20, cells were stained with the anti-ALPP/L2-PE antibody (Merck). The top and bottom 10% cells were sorted using a BD Aria sorter (BD biosciences). Then, DNA was extracted using the QIAwave DNA kit (Qiagen), according to the manufacturer’s protocol. The genomic DNA was amplified by PCR using the following primer sequences: P5: AATGATACGGCGACCACCGAGATCTACACTCTTTCCCTACACGACGCTCTTCCGATCT[s]TTGTGGAAAGGACGAAACACCG. P7: CAAGCAGAAGACGGCATACGAGATCGGTTCAAGTGACTGGAGTTCAGACGTGTGCTCTTCCGATCTTCTACTATTCTTTCCCCTGCACTGT. Then, PCR products were purified using QIAquick PCR purification kit (Qiagen) according to the manufacturer’s protocol.

Samples were sequenced by a third-party vendor (Azenta); the resulting fastq raw data were processed into count matrices using PoolQ. QC (data not shown) confirmed sufficient guide recovery (>98% for sgRNAs with count >20) and even guide count distributions. Performance of controls was assessed by comparing the samples to PlasmidDNA: the data showed a sufficient decrease in signal in positive controls (common essential genes) and negative control distributions (non-targeting, intergenic targeting, and common non-essential genes), closely aligned with the bulk of the non-control targets in the data set. The source of essential and non-essential genes considered in this analysis was Hart et al ^47^. For gene-level analysis, the data were scored using MAGECK RRA ^48^ as well as the hypergeometric algorithm (https://github.com/mhegde/volcano_plots) by comparing the ALPG-high to ALPG-low and ALPP-high to ALPP-low samples for ALPP-KO and ALPG-KO, respectively. For the hypergeometric analysis, inputs were gene-level log2-fold changes calculated as averaged guide-level log2-fold changes based on log2-count-per-million normalized guide counts. Hits were identified using a consensus approach considering genes that passed a p-value < 0.05 across both algorithms. Genes targeted by fewer than 3 or more than 10 sgRNAs were excluded from the analysis.

### Overrepresentation Analysis

The overrepresentation analysis was done using a Fisher’s exact test, testing positive regulator hits against gene sets from the molecular signature database (MSigDb). Canonical pathways (C2:CP) and gene ontology sets (C5:BP, biological pathways; C5:MF, molecular functions) were included as sets of interest, and the number of non-control targets in the genome-wide library was used as background size. P values from Fisher’s exact test were FDR-corrected for multiple hypothesis testing. Results were filtered based on gene set size (<350 genes) and FDR-threshold (<0.01).

### Capillary-based Western Blot

Whole-cell protein extracts in RIPA buffer supplemented with 1x protease/phosphatase inhibitors (Invitrogen) were diluted to 0.16 mg/mL in 0.1X Sample Buffer (Biotechne (042-195)) containing 1X Fluorescent Master Mix (Biotechne (PS-FL01-8) and incubated for 15 minutes at 60°C in a Thermal Cycler (Bio-Rad (1861096)) for denaturation. Then, the proteins were loaded on a 12- 230 kDa Separation Module (Biotechne (SM-W001)) at 3 µL per well (Row A). Primary antibodies against NF2 (Cell Signaling (6995); 1:20), TEAD4 (Abcam (ab58310); 1:25), ALPP/ALPPL2 (Cell Signaling (30971); 1:30), and Beta-Actin (Cell Signaling (4970); 1:50) were diluted in Antibody Diluent 2 (Biotechne (042-203)) and loaded at 10 µL per well (Row C). Pre-diluted Anti-Rabbit Secondary HRP Antibody (Biotechne (042-206)) and Anti-Mouse Secondary HRP Antibody (Biotechne (042-205)) were loaded at 10 µL per well (Row D). Luminol-S (Biotechne (043-311)) and Peroxide (Biotechne (043-379)) were mixed at a 1:1 ratio and loaded at 15 µL per well (Row E). 500 µL of Wash Buffer (Biotechne (042-202)) was distributed in the corresponding wells. After centrifugation at 1,000 xG for 5 minutes, the filled plate is loaded on a Jess Automated Western Blot System (Biotechne (004-650)), along with the Capillary Cartridges (Biotechne (PS- CC01)). After running, the results are analyzed on Compass for Simple Western software (Biotechne).

### *In vitro* treatments

3.104 cells were treated with the indicated concentration of Pan-TEADi (MedChemExpress), Kras G12Ci (Cayman Chemical), Pan-Rasi (MedChemExpress), MEKi (Cayman Chemical), and ERKi (Cayman Chemical) for three days before evaluation of ALPP/L2 expression by flow cytometry.

G12Ci resistant cell lines were generated by subsequential increase of G12Ci inhibitor concentration starting at IC10 over a 6-month time period, with 2-3 doublings per concentration until 100x IC50 was reached.

### Antibody Drug Conjugate Cytotoxicity assay

SW837 cell lines were pre-treated with 62.5 nM of Kras G12Ci (Cayman Chemical) or DMSO (Sigma-Aldrich) for 48 hours. Then, the cells were washed twice to remove dead cells and Kras G12Ci. 2.10^3^ pre-treated or untreated live SW837 WT, SW837 OE, SW837 DKO, SW837 TEAD4 KO, SW837 NF2 KO, NCI-H358 DMSO or NCI-H358R cells were incubated with various concentrations of ALPP/L2-MMAE (Merck) or isotype-MMAE (Merck) conjugates for 6 days. Cells were incubated 30min with 1:1 (v/v) of CellTiter-Glo Luminescent viability assay (Promega). Cell viability was normalized to untreated groups.

### RNA extraction and real-time PCR for gene expression

Total RNA was isolated using the MagMAX mirVana Total RNA isolation kit (A27828, Thermo Fisher Scientific Inc., Foster City, CA) according to the manufacturer’s instructions. Total RNA samples were qualified and quantified on the Fragment Analyzer (Agilent, Santa Clara, CA) per manufacturer’s instructions. DNase-treated total RNA was reverse-transcribed using QuantiTect Reverse Transcription (Qiagen, Valencia, CA) according to manufacturer’s instructions. 20X primer assays were obtained commercially from Thermo Fisher Scientific (Foster City, CA). Gene specific pre-amplification was done on 10 ng cDNA per Standard BioTools Biomark manufacturer’s instructions (Standard BioTools, Foster City). Real-time quantitative PCR was then done on the Standard BioTools Biomark HD using 20X Taqman primer assays (Thermo Fisher Scientific Inc, Foster City, CA) with Taqman Fast Universal PCR Master Mix with no AmpErase UNG. Samples and primers were run on either a 48.48 Dynamic Array or 96.96 Dynamic Array or 192.24 Dynamic Array per manufacturer’s instructions (Standard BioTools, Foster City). UbiquitinB levels were measured in a separate reaction and used to normalize the data by the Δ Ct method. (Using the mean cycle threshold value for ubiquitin and the gene of interest for each sample, the equation 1.8 ^ (Ct UbiquitinB minus Ct gene of interest) x 104 was used to obtain the normalized values.)

### Animal studies Mice

Wild-type C57BL/6NTac, ADORA2A KO mice were obtained from Taconic. C57BL/6J mice were obtained from Jackson Laboratories. Mice were maintained under specific pathogen-free conditions and kept in microisolators with filtered air at the research laboratories of Merck & Co., Inc., Rahway, NJ, USA (MRL) animal facilities. All animal procedures were approved by the South San Francisco Institutional Animal Care and Use Committee in accordance with guidelines of the Association for Assessment and Accreditation of Laboratory Animal Care.

### Cell line-Derived Xenograft (CDX) models

7-9 weeks C57BL/6NTac, ADORA2A KO, C57BL/6J mice were implanted subcutaneously with MC38 EV, MC38 hALPPL2WT, MC38 hALPPL2 S111A, MB49 EV or MB49 hALPPL2WT cells. At day 19 (MB49) or day 21 (MC38), tumors were collected, dissociated, and stained for flow cytometry analysis.

8-week-old female NSG mice (NOD.Cg-*Prkdc^scid^ Il2rg^tm1Wjl^*/SzJ) (The Jackson Laboratories) were implanted subcutaneously with SW837 cells. When tumors reached 150-200mm3, mice were randomized and dosed p.o. with a 10mpk of Kras G12Ci (Cayman Chemical) or vehicle control (HPMC 2% (Sigma-Aldrich), Tween 80% (Sigma-Aldrich) 1%, water 97% (v/w)) every day for a total of eight doses. At day 2 post Kras G12Ci treatment, mice were dosed intravenously with 3mpk of ALPP/L2-MMAE or isotype-MMAE. For evaluation of ALPP/L2 expression by flow cytometry, tumors were collected at day 2 post-Kras G12Ci treatment, dissociated, and stained.

### ALPP homology model generation and ATP docking

Homology models were created using UniProtKB sequence ID P10696 and RCSB ID:1EW2 as a template using the Homology Modeler application in MOE 2022.02 (http://www.chemcomp.com/index.htm (2022)). A 0.5 kcal/mol gradient limit was used during refinement. The structure was aligned and overlaid with the template, retaining metal ions and solvent. ATP was docked to sites found in close proximity to metal ions, and structures were minimized with the Amber14:EHT forcefield ^49,50^

### Recombinant ALPP and ALPPL2 cloning and purification

Human ALPP (uniprot:P05187) and ALPPL2 or ALPG (uniprot:P10696) are 96% identical. For ALPP aa 23-507 and ALPPL2 aa 20-503 were chosen as boundaries for recombinant expression. Cloning included SLAM leader peptide at the N-termini of the coding sequence that directs secretion of the recombinantly expressed protein into the surrounding media, and at the C-termini was a 6x histidine tag to enable affinity purification using Ni-charged resins. The coding sequences were cloned under a constitutively active CMV promoter in a pTT5-derived mammalian expression vector. Different point mutants were made to both ALPP and ALPPL2 to effect changes to enzyme activity. For ALPP, S114A, R188M, H454I, H175I, D113N, D379V, S114A/R188M/H454I/H175I, D113N/D379V and S114A/R188M/H454I/H175I/D113N/D379V mutants were made alongside the wildtype (WT) enzyme. For ALPPL2, S111A, R185M, H451I, H172I, D110N, D376V, S11A/R185M/H451I/H172I, D110N/D376N, and S111A/R185M/H451I/H172I/D110N/D376N mutants were designed and made alongside the WT molecule.

Transient expression of these designed molecules involved chemical transfection enabled by PEIMax pH 7.0 into Horizon Discovery CHO cells (HD-CHO) at a cell density of 6 million cells/mL. The cells were cultured in tubespin bioreactors at 25 mL culture volume, grown at 36.5°C, 5% CO2, 80% relative humidity, with a shake speed of 300 rpm, using a Kuhner incubator with a 25 mm throw. The temperature was changed to 33°C after feed on day 1, with another feed on day 5. Cell culture was harvested on day 7 with viabilities >80% and clarified by centrifugation/filtration with 0.22 μm membrane filters before proceeding to affinity purification.

The purification of these constructs came from a one-step affinity purification of 25 mL clarified supernatant with Ni sepharose excel (Cytiva) resin with a settled volume of 0.2 mL as a 50% slurry in PBS (phosphate-buffered saline) pH 7.4. The protein was eluted in 4CV (column volumes) and buffer-exchanged into a final formulation of 10 mM sodium phosphate, 75 mM sodium chloride, 3% sucrose, pH 7.4 buffer for further analysis and –80°C storage.

### Malachite green phosphate assay

10 ng of recombinant ALPP or ALPPL2 proteins (Genscript) or ALPPL2-expressing MC38 cells were incubated for 30 minutes at 37°C with different concentrations of ATP, ADP, and AMP (Sigma-Aldrich) in 1X NE buffer 2 (New England Biolabs). After 30 minutes, the malachite green phosphate reagent (Sigma-Aldrich) was added and incubated for 30 minutes at room temperature. The absorbance at 620nm was measured on the plate reader SpectraMax I3x (Molecular Device).

### Adenosine assay

100 ng of recombinant ALPPL2 WT protein (Genscript) was incubated for 25 minutes at 37°C with 10 nM of ATP (Sigma-Aldrich) in Alkaline phosphatase assay buffer (Abcam). 1:1 (v/v) adenosine reaction mix from the adenosine assay kit (Abcam), according to the manufacturer’s protocol. Incubate 15 minutes at room temperature in the dark. The fluorescence signal (Ex/Em=535/587nm) was measured on the plate reader SpectraMax I3x (Molecular Device).

### Protocol antibody conjugation and QC

To a solution of the ALPP/L2 or isotype control antibody (Merck) (5.00 mg, 0.033 µmol) was added an aqueous solution of 50 mM Tris(2-carboxyethyl)phosphine hydrochloride (20 eq., 13.4 µL, 0.67 µmol), and the mAb was reduced for 2 h at 37 °C. The reduced antibody was exchanged into 1x PBS buffer (pH 7.4) and diluted to a concentration of 10 mg/mL in 1x PBS buffer (500 µL). To the antibody solution was slowly added a 10 mM solution of the linker-payload (4 eq.) in DMSO, with vigorous mixing. The reaction was then agitated at RT for 16 h. The ADC was purified by SEC (AKTA Pure*TM,* Cytiva Superdex 200 Increase 10/300 GL, 1x PBS pH 7.4 buffer, monitoring at 280 nm) and was characterized by LC-MS (Agilent 1290 Infinity II, Agilent PLRP-S, 1000 Å, 5 µm, 2.1 x 50mm, 25-70% MeCN/H_2_O with 0.1% TFA, 80°C column temperature) and SEC (Agilent 1290 Infinity II, Waters Acquity UPLC Protein BEH SEC, 200 Å, 1.7 µm, 4.6 x 150 mm, 100 mM sodium phosphate, 200 mM NaCl, 5% IPA, 30°C column temperature).

### Determination of Drug-to-Antibody Ratio (DAR)

The drug-to-antibody ratio was determined using the reduced antibody drug conjugate masses of the heavy and light chains. The antibody drug conjugate was reduced with Tris(2- carboxyethyl)phosphine hydrochloride and analyzed by reversed-phase chromatography with mass spectrometry on an Agilent 1290 Infinity II instrument with an Agilent AdvanceBio 6545XT LC/Q-TOF detector. The running buffer was MeCN/H_2_O with 0.1% TFA. The observed signal peaks for the reduced antibody drug conjugate were deconvoluted using Agilent Bioconfirm Software. The individual drug loading peaks of the heavy and light chains and intensities were used to determine the drug-to-antibody ratio of the individual chains and combined to determine the overall drug-to-antibody ratio of the antibody drug conjugate.

### ATAC-seq

ATAC-seq library preparation and sequencing reactions were conducted at Azenta Life Sciences (South Plainfield, NJ, USA). Live cell samples were thawed, washed, and treated with DNAse I (Life Tech, Cat. #EN0521) to remove genomic DNA contamination. Live cell samples were quantified and assessed for viability using a Countess Automated Cell Counter (ThermoFisher Scientific, Waltham, MA, USA). After cell lysis and cytosol removal, nuclei were treated with Tn5 enzyme (Illumina, Cat. #20034197) for 30 minutes at 37°C and purified with Minelute PCR Purification Kit (Qiagen, Cat. #28004) to produce tagmented DNA samples. Tagmented DNA was barcoded with Nextera Index Kit v2 (Illumina, Cat. #FC-131-2001) and amplified via PCR prior to a SPRI Bead cleanup to yield purified DNA libraries.

The sequencing libraries were multiplexed and clustered onto a flowcell on the Illumina NovaSeq instrument according to the manufacturer’s instructions. The samples were sequenced using a 2x150bp Paired End (PE) configuration. Image analysis and base calling were conducted by the NovaSeq Control Software (NCS). Raw sequence data (.bcl files) generated from Illumina NovaSeq were converted into fastq files and de-multiplexed using Illumina bcl2fastq 2.20 software. One mismatch was allowed for index sequence identification.

ATAC-seq data generated in this study were preprocessed and mapped following Galaxy training materials. Filtered and cleaned BAM files were converted to BED files using bedtools genomecov using the –bg option. BED files were then converted to bigWig format using the UCSC bedGraphtoBigWig tool and visualized using IGV 2.19.5 (https://trainings.galaxyproject.org, ^51,52^) PC9 ATAC-seq data were obtained from the ChIP-atlas (https://chip-atlas.org/, accession number SRX19752611).

### ChIP-seq

ChIP-seq data were downloaded from the ChIP-atlas (https://chip-atlas.org/). PC9 DMSO a: TEAD4 (SRX5887617); PC9 a: YAP1 (SRX5887621); PC9 input control (SRX1425832); SNU216 a: TEAD4 (SRX243629); SNU216 Input Ctrl (SRX243628); HepG2 a: TEAD4 (SRX190331); HepG2 Input Ctrl (SRX5574517)

### Statistics

One-way ANOVA, unpaired t-test and paired t-test were used to calculate statistical significance in the rest of this study. Ns, Not Significant, * p<0.05, ** p<0.01, *** p<0.001, **** p<0.0001. Statistics were performed using GraphPad Prism 7 software.

## DECLARATIONS

### Ethics approval

All animal procedures were approved by the Merck & Co., Inc., Rahway, NJ, USA Animal Care and Use Committee in accordance with guidelines of the Association for Assessment and Accreditation of Laboratory Animal Care.

## Acknowledgments

We would like to acknowledge our internal colleagues Codi Convington, Ruban Mangadu, Danye Cheng and Randolph Smith, Debajan Barua for their technical support, Su-Yang Liu for helpful discussions.

## Author Contributions

Conceptualization: J.L.M; B.H; D.B, Investigation and Validation: J.L.M; B.H; T.E; A.R.H.; M.D; R.P; W.H.K; S.M; C.L; E.M; K.P; K.N.; S.H; K.S; P.Y; D.B, Formal analysis: J.L.M; B.H; T.E; A.R.H.; M.D; R.P; W.H.K; E.M; S.S.W.H; P.Y D.B, Methodology: J.L.M; B.H; W.H.K; S.H; E.M; K.S; D.B., Data Curation: T.E; A.R.H; M.D, Project administration: D.B Resources: B.R Supervision: O.O; S.S, D.B, Visualization: J.L.M; B.H; D.B, Writing-original draft: D.B

