## Supplemental figures for "YAP/TEAD4-regulated placental alkaline phosphatases ALPP and ALPPL2 are immunosuppressive ectonucleotidases modulated by MAPK inhibitors"

### Figure S1

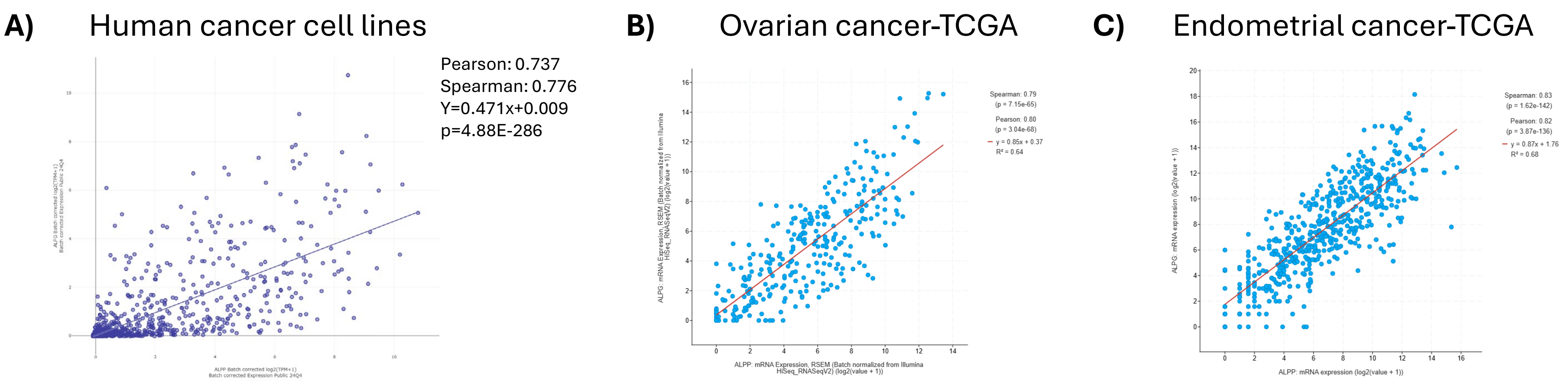


**Figure S1: ALPP and ALPPL2 expression in normal tissues and tumor cells**

Positive correlation of ALPP and ALPG expression in human cancer cell lines from DepMap[53] **(A)**, human primary tumor cells from Ovarian (n=489) **(B),** or Endometrial (n=548) **(C)** patients. [54,55]

### Figure S2

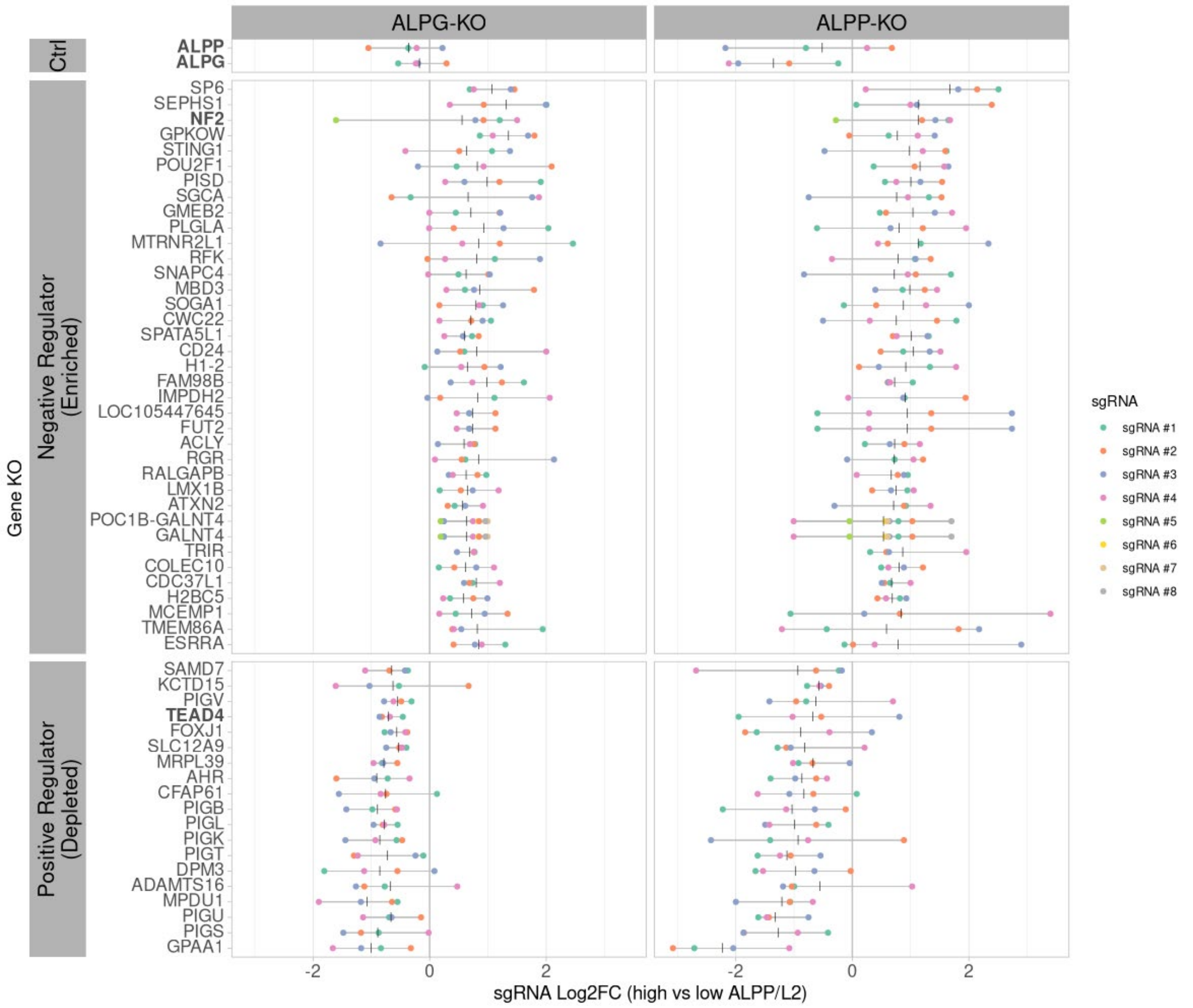


**Figure S2: Significant negative and positive regulators of ALPP/L2 in tumor cells**

Fold change of the significant hits in ALPGKO- or ALPPKO-high versus ALPGKO- or ALPPKO-low cells from individual sgRNAs. Colors distinguish the different sgRNAs for each gene to enable comparison across screens; vertical bars indicate averages across sgRNAs.

### Figure S3

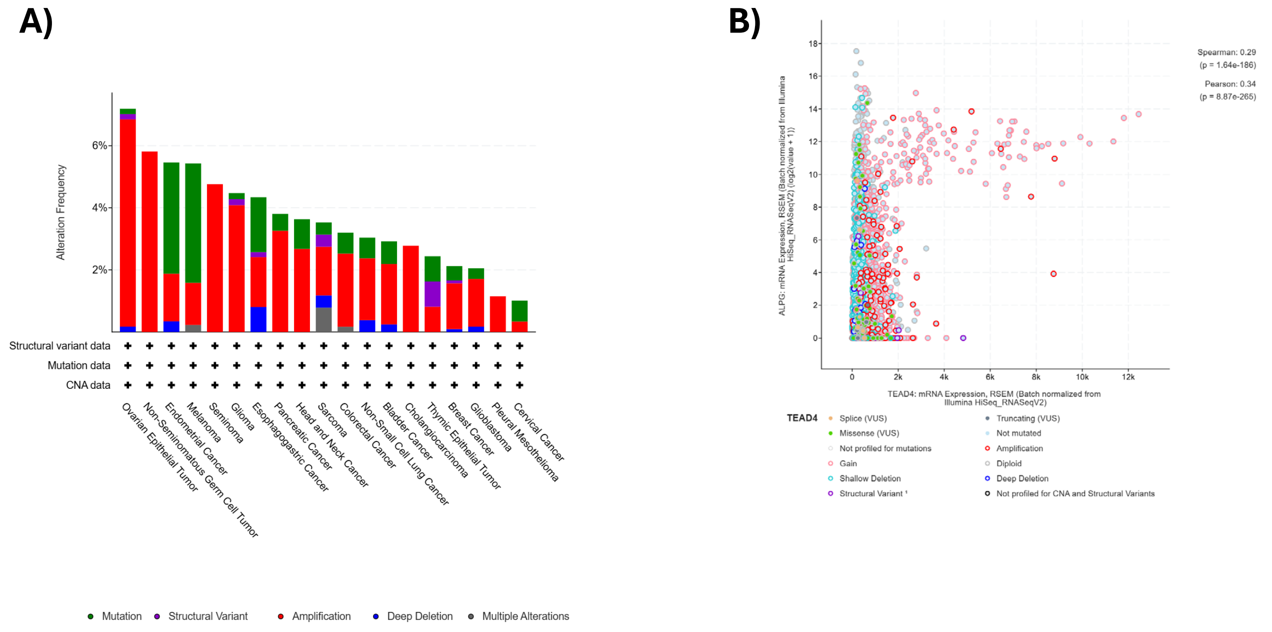


**Figure S3: TEAD4 is amplified in ALPP/L2 positive tumors**

**(A)** TEAD4 alteration in various cancer types. TEAD4 is commonly amplified in ovarian, testicular, PDAC, endometrial, and NSCLC. **(B)** TEAD4 and ALPG co-expression in cancer samples from multiple indications. High TEAD4 expression is correlated with ALPG high expression.

### Figure S4

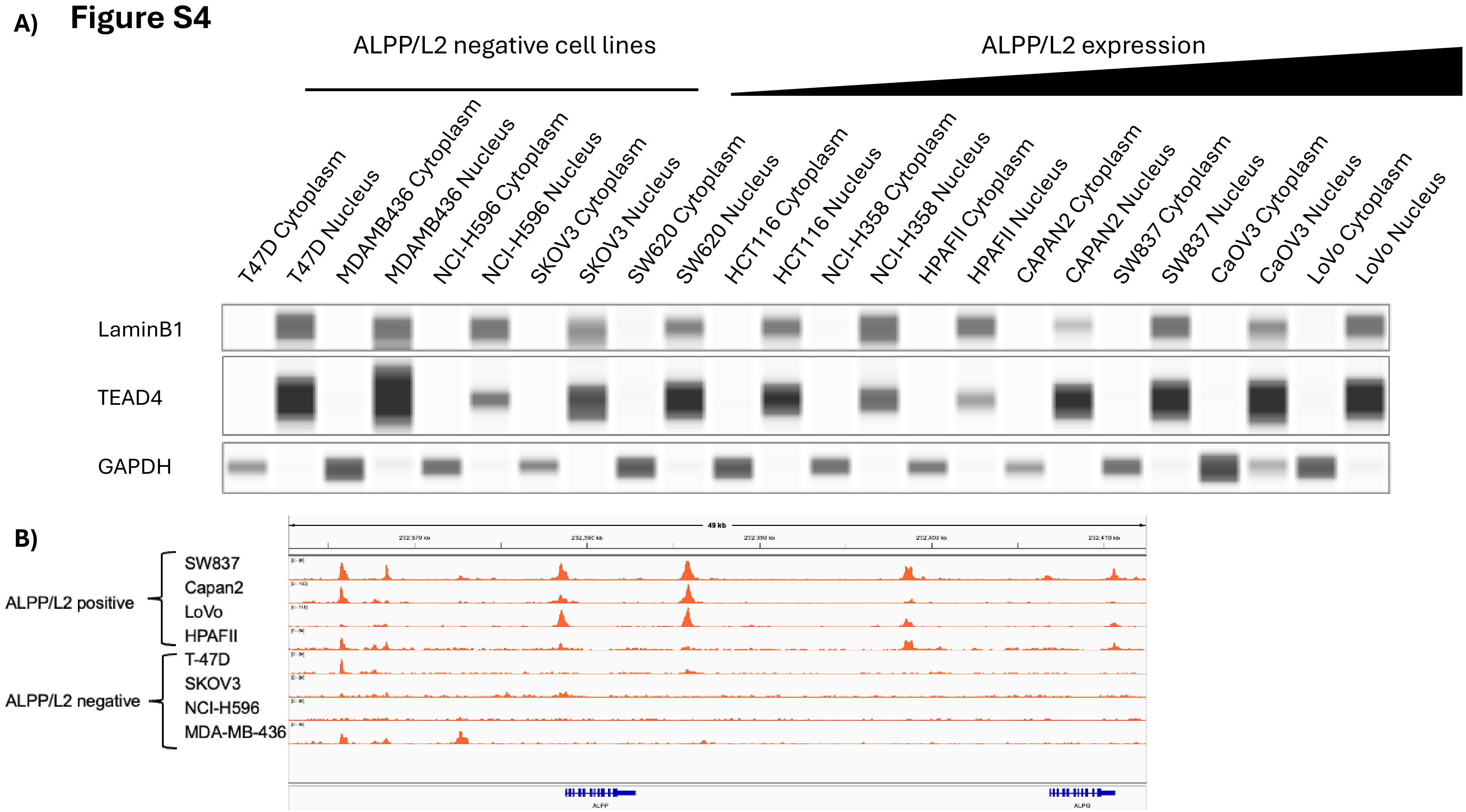


**Figure S4: ALPP/L2 expression correlates with chromatin accessibility but not TEAD4 subcellular localization**

**(A)** TEAD4 expression in the cytoplasmic- as shown by GAPDH staining- or nuclear -as shown by LaminB1 staining- fractions of human cancer cell lines expressing different levels of expression of ALPP/L2. T47D, MDA-MB-436, NCI-H596, and SKOV3 cells are negative for ALPP/L2. **(B)** ATAC-seq data from ALPP/L2 negative and positive cell lines showing open chromatin regions in ALPP and ALPPL2 promoters.

### Figure S5

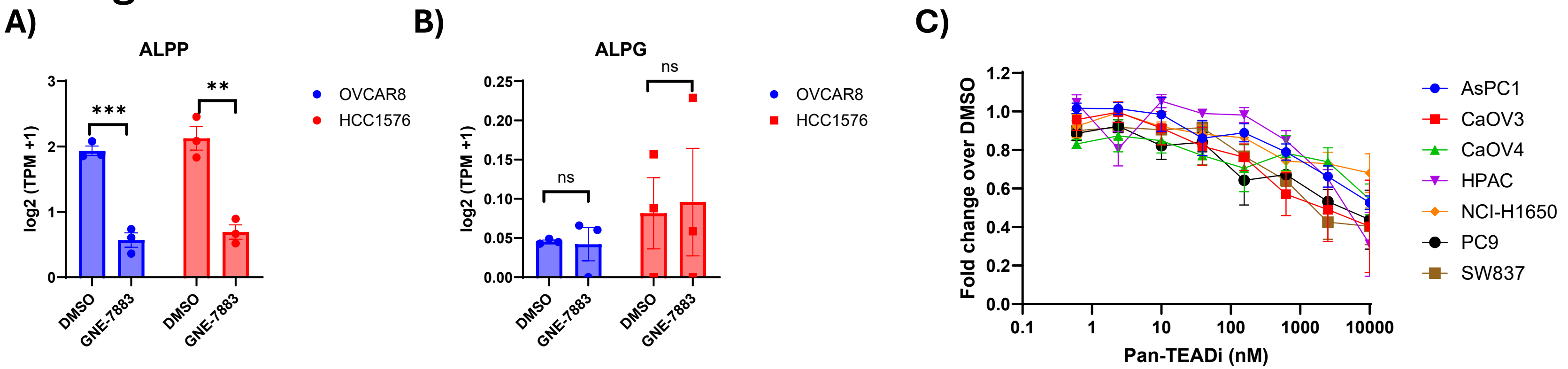


**Figure S5: Pan-TEAD inhibitors inhibit ALPP/L2 expression in human cancer cell lines**

**(A)** *ALPP* and **(B)** *ALPG* expression from OVCAR8 and HCC1576 cells treated for 48 hours with DMSO or GNE-7883 was extracted from published RNA-seq (GSE229066 [13]). **(C)** ALPP/L2 expression was evaluated by flow cytometry on the surface of ALPP/L2-expressing human cancer cell lines 72 hours post-treatment with DMSO or Pan-TEADi. Data are represented as mean ± SEM. n=3-5 independent experiments.

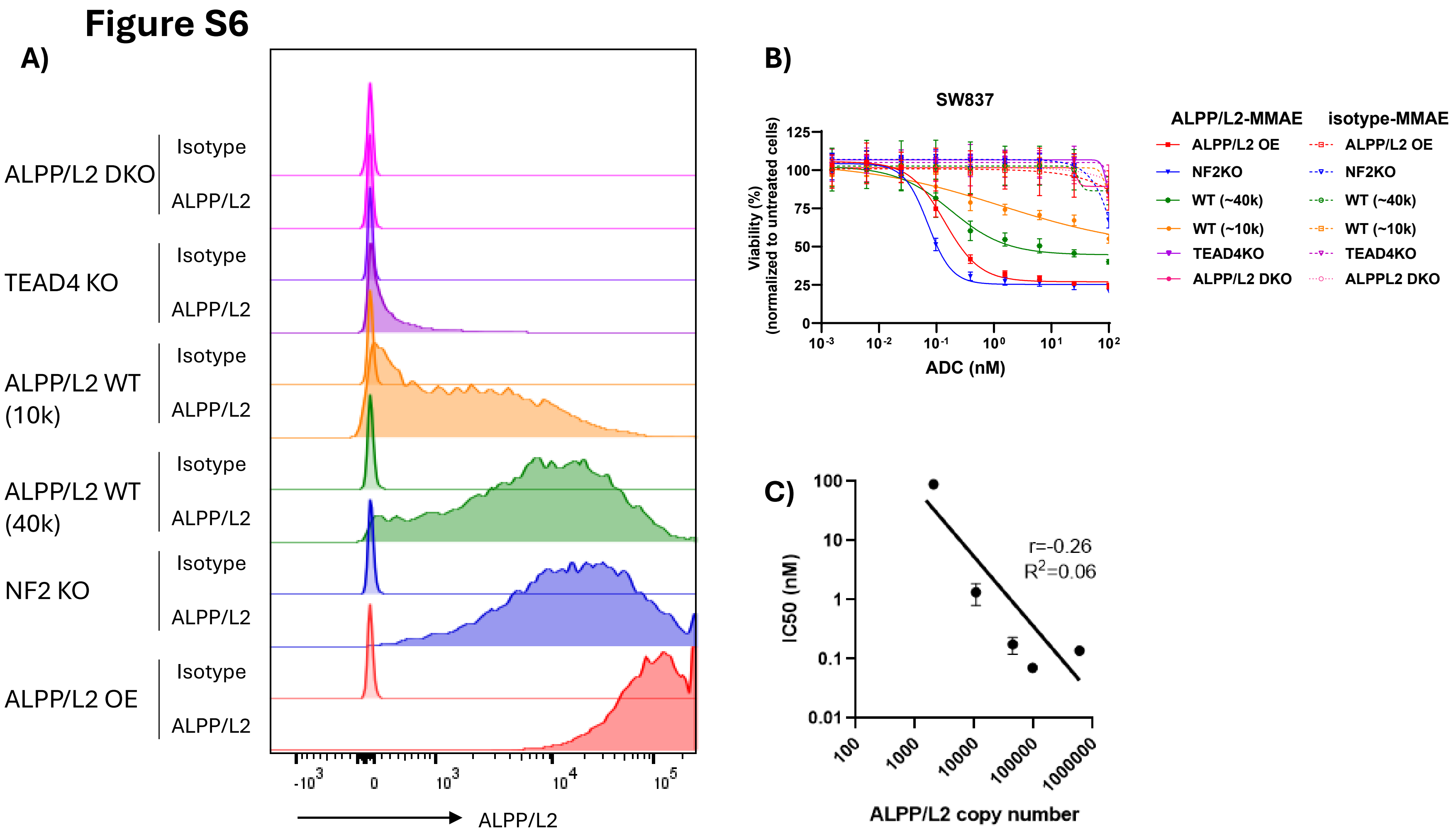


**Figure S6: ALPP/L2 surface expression correlates with ALPP/L2-MMAE potency *in vitro***

**(A)** Representative histogram of ALPP/L2 expression evaluated by flow cytometry on the surface of SW837 ALPP/L2 DKO, SW837 WT (10k or 40k), SW837 ALPP/L2 over-expressing (OE), SW837 TEAD4 KO, or SW837 NF2 KO cells. **(B)** Cell viability of the listed SW837 cells treated with different concentrations of ALPP/L2-MMAE or Isotype-MMAE conjugates for 6 days. **(C)** Pearson correlation of ALPP/L2 surface expression with ALPP/L2-MMAE potency *in vitro,* as shown by the IC50 values.

Data are represented as mean ± SEM. n=2 independent experiments.

### Figure S7

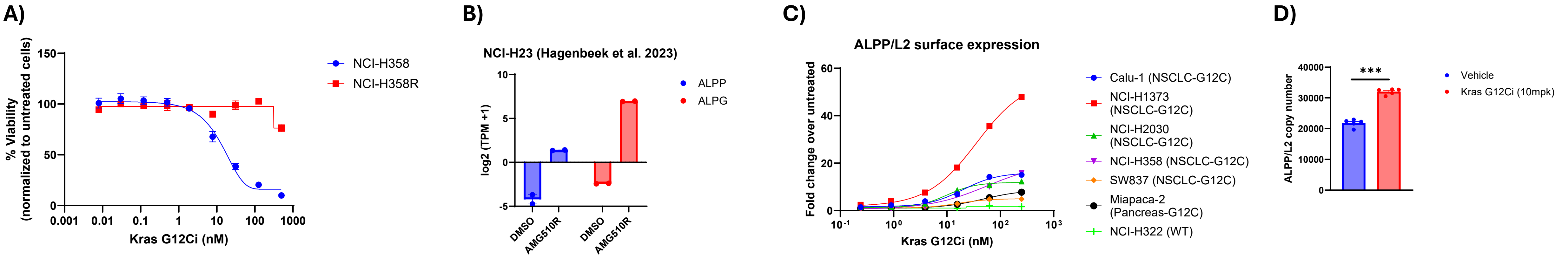


**Figure S7: Kras G12Ci treatment induces ALPP/L2 expression in human cancer cell lines *in vitro* and *in vivo***

**(A)** Viability of NCI-H358 DMSO or NCI-H358R treated with various concentrations of Kras G12Ci for three days. **(B)** *ALPP* and *ALPG* expression in NCI-H23 AMG510R compared to DMSO-treated cells (DMSO), measured by RNA-seq (GSE229066 [13]). **(C)** Dose-dependent upregulation of ALPP/L2 on the surface of Kras G12C mutant human cancer cell lines. Data shown as fold change of ALPP/L2 copy number over DMSO control. Representative histogram and quantification of ALPP/L2 cell surface expression on NCI-H358 DMSO or NCI-H358R cells *in vitro*. **(D)** *ex-vivo* ALPP/L2 copy number at the surface of SW837 tumor cells, 2 days post treatment with Kras G12Ci at 1 or 10mpk.

Data are represented as mean ± SEM. **(A-C)** n=2-3, **(D)**n=1 independent experiments. Unpaired t-test ns=non-significant, * p<0.05, *** p<0.001

### Figure S8

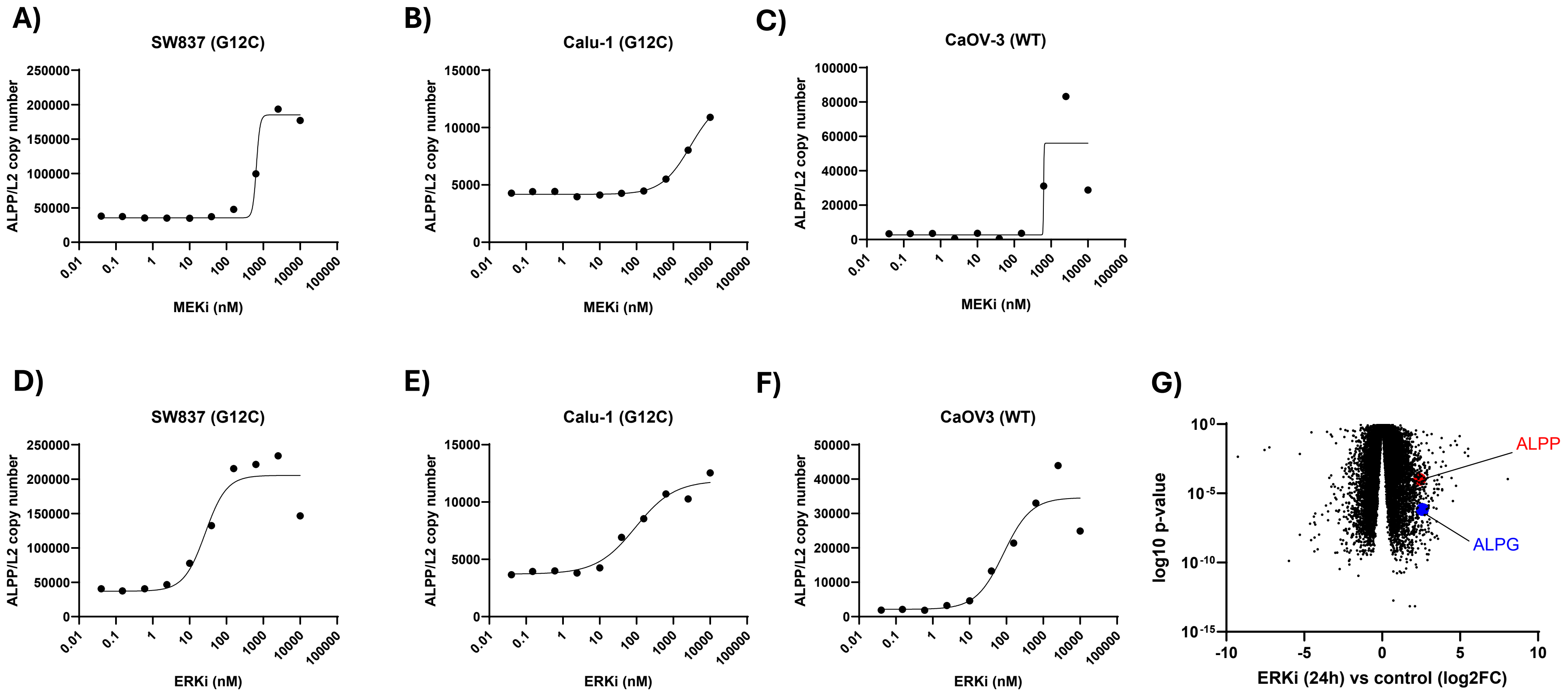


**Figure S8: MEK and ERK inhibitors treatment induces ALPP/L2 expression in human cancer cell lines**

**(A-C)** Dose-dependent upregulation of cell surface ALPP/L2 on SW837 cells **(A)**, Calu-1 **(B)**, or CaOV3 **(C)** treated with MEKi. **(D-F)** Dose-dependent upregulation of cell surface ALPP/L2 on SW837 cells **(D)**, Calu-1 **(E)**, or CaOV3 **(F)** treated with MEKi. **(G)** Differential gene expression analysis for pancreatic cell lines treated with SCH772984 (ERK inhibitor) at 1 μM for 24 hours compared to paired untreated control cells (PRJEB25806, [56]) ALPP and ALPG are highlighted in red and blue, respectively.

Data are representative of **(A-F)** n=3 independent experiments.

### Figure S9

A) B) C)

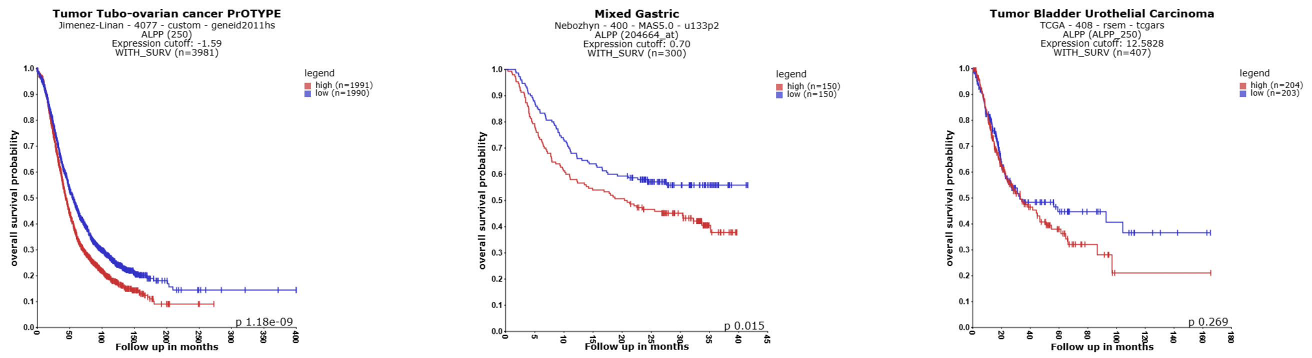


**Figure S9: High ALPP expression is associated with poor prognosis in multiple tumor types**

Kaplan-Meier curves of ALPP in **(A)** Ovarian cancer (n=1990- 1991 patients) Jimenez-Linan (GSE135820), **(B)** Mixed Gastric cancer (n=150 patients) (GSE66229), **(C)** Bladder cancer (n=203-204 patients) (TCGA). Cutoff median expression. Logrank test as described [57]. Data generated from R2: Genomics Analysis and Visualization Platform (http://r2.amc.nl).

### Figure S10

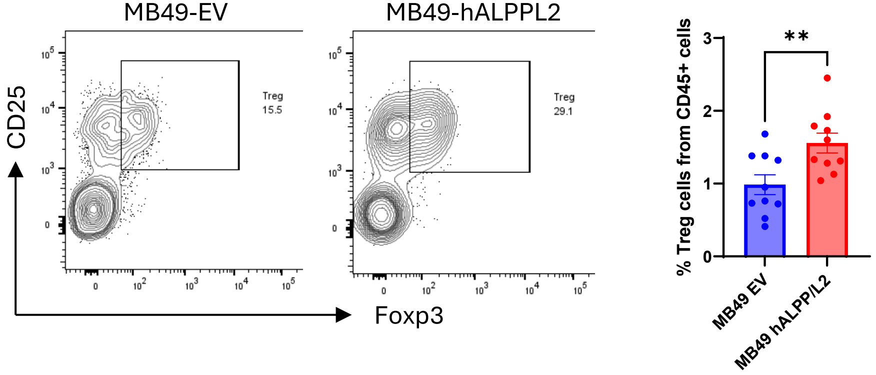


**Figure S10: ALPPL2 expression promotes Treg infiltration in MB49 syngeneic tumor model.**

Representative flow cytometric analysis of Treg cells (live CD45+CD3+CD4+Foxp3+CD25+) (left) and quantification (right) in MB49-EV and MB49-hALPPL2 tumors, day 19 post-implant. (n=10 mice per group)

### Figure S11

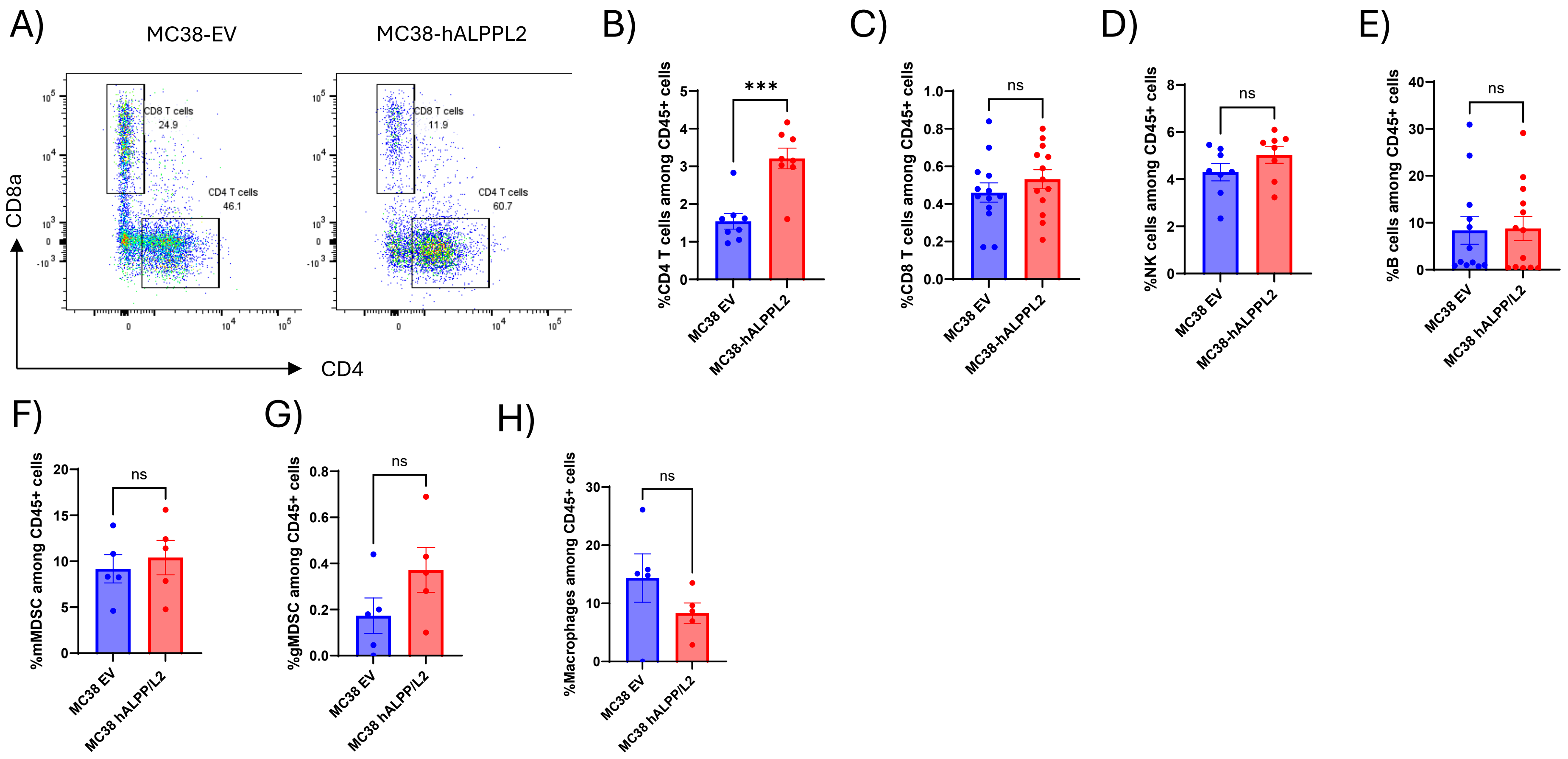


**Figure S11: Immunophenotyping of MC38 EV and MC38-hALPPL2 tumors in C57BL/6 mice**

**(A)** Representative flow cytometric analysis of tumor-infiltrating CD4 and CD8 T cells (gated on live CD45+CD3+NK1.1-). **(B-H)** Quantification of the proportion of **(B)** CD4 T cells (CD45+CD3+NK1.1-CD4+CD8a-), **(C)** CD8 T cells (CD45+CD3+NK1.1-CD4-CD8a+), **(D)** NK cells (CD45+CD3-NK1.1+), **(E)** B cells (CD45+CD3-NK1.1-CD19+), **(F)** mMDSC cells (CD45+CD11bhigh Ly6Chigh Ly6G low), **(G)** gMDSC cells (CD45+CD11bhigh Ly6Clow Ly6Ghigh), **(H)** macrophage cells (CD45+CD11b+CD11c+F4/80+) among live CD45+ cells at day 21 post implant. Data are represented as mean ± SEM. n=2-3 independent experiments. Unpaired t-test ns=non-significant, *** $p<0.001$

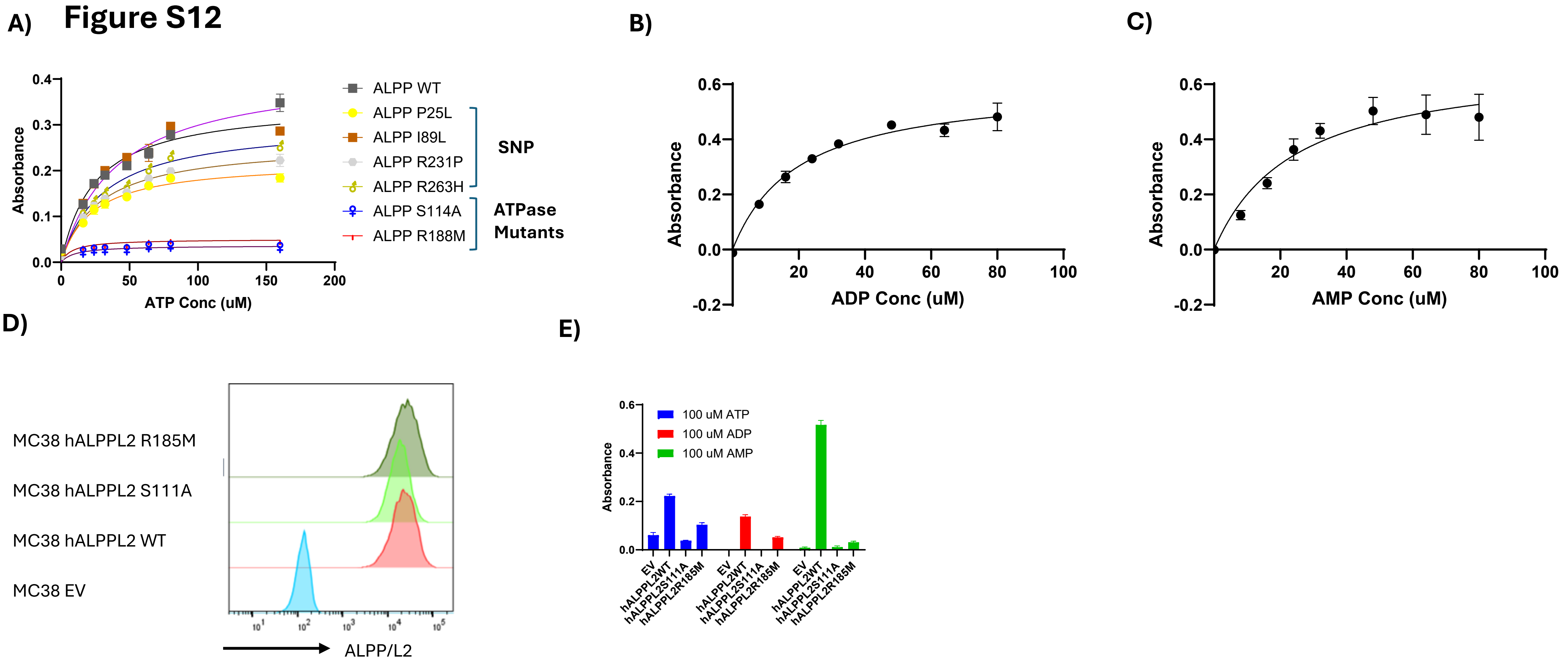


**Figure S12: S111A mutation abrogates ALPP and ALPPL2 ectonucleotidase function**

**(A)** ATPase activity of ALPP WT, SNPs, S114A, and R188M mutants using malachite green assay Dose-dependent hydrolyzation of **(B)** ADP and **(C)** AMP by recombinant human ALPP. **(D)** Representative histogram of ALPP/L2 cell surface expression on MC38 EV, MC38 hALPPL2 WT, MC38 hALPPL2 S111A, and MC38 hALPPL2 R185M cells. **(E)** Phosphatase activity of MC38 EV, MC38 hALPPL2 WT, MC38 hALPPL2 S111A, and MC38 hALPPL2 R185M cells treated with ATP, ADP, or AMP.
